# Silent Crickets Reveal the Genomic Footprint of Recent Adaptive Trait Loss

**DOI:** 10.1101/489526

**Authors:** Sonia Pascoal, Judith E. Risse, Xiao Zhang, Mark Blaxter, Timothee Cezard, Richard J. Challis, Karim Gharbi, John Hunt, Sujai Kumar, Emma Langan, Xuan Liu, Jack G. Rayner, Michael G. Ritchie, Basten L. Snoek, Urmi Trivedi, Nathan W. Bailey

**Author notes:** Contributed equally (Sonia Pascoal, Judith E. Risse, Xiao Zhang).

## Abstract

Secondary trait loss is widespread and has profound consequences, from generating diversity to driving adaptation. Sexual trait loss is particularly common^1^. Its genomic impact is challenging to reconstruct because most reversals occurred in the distant evolutionary past and must be inferred indirectly^2^, and questions remain about the extent of disruption caused by pleiotropy, altered gene expression and loss of homeostasis^3^. We tested the genomic signature of recent sexual signal loss in Hawaiian field crickets, *Teleogryllus oceanicus*. Song loss is controlled by a sex-linked Mendelian locus, *flatwing*, which feminises male wings by erasing sound-producing veins. This variant spread rapidly under pressure from an eavesdropping parasitoid fly. We sequenced, assembled and annotated the *T. oceanicus* genome, produced a high-density linkage map, and localised *flatwing* on the X chromosome. We characterised pleiotropic effects of *flatwing*, including changes in embryonic gene expression and alteration of another sexual signal, chemical pheromones. Song loss is associated with pleiotropy, hitchhiking and genome-wide regulatory disruption which feminises flatwing male pheromones. The footprint of recent adaptive trait loss illustrates R. A. Fisher’s influential prediction that variants with large mutational effect sizes can invade genomes during the earliest stages of adaptation to extreme pressures, despite having severely disruptive genomic consequences.

---

Male crickets sing to attract and court females and to fight with rivals, but approximately 15 years ago, silent *T. oceanicus* males arose and spread in populations on the Hawaiian archipelago^4,5^ (Fig. 1a). They were first detected in 2003 in a population on Kauai, where they rapidly spread to near-fixation from undetectable starting frequencies, under selection imposed by a lethal parasitoid fly, *Ormia ochracea* (Fig. 1b)^4^. Female flies acoustically locate male crickets by eavesdropping on their songs, but silent flatwing males have feminised wings lacking structures used to produce sound and are thus protected (Fig. 1c). The genetic mutation(s) underlying the flatwing phenotype show Mendelian segregation and X-linkage^6,7^, and the propagation of flatwing males to near-fixation in the Kauai population represents one of the fastest rates of evolution known in the wild, having occurred in fewer than 20 generations^4^. All males found in a comprehensive survey of this population in October 2018 were flatwing (38 flatwing males, no normal-wing males found or heard singing by JGR and NWB), but the continued existence of the population indicates that silent males still find mates and must compensate for their inability to sing. The selective environment promoting the rapid spread of flatwing crickets is understood, but the mechanistic basis of the phenotype remain an open question. How did such a spectacularly disruptive phenotypic change invade the genome of crickets so quickly? Foundational evolutionary theory predicts that adaptive variants which invade genomes and spread under positive selection should tend to be small in effect size and exert few pleiotropic consequences, although exceptions are predicted during the earliest stages of adaptation^8,9^. Empirical studies have been unable to address this in naturally-evolving systems.

**Fig. 1.**
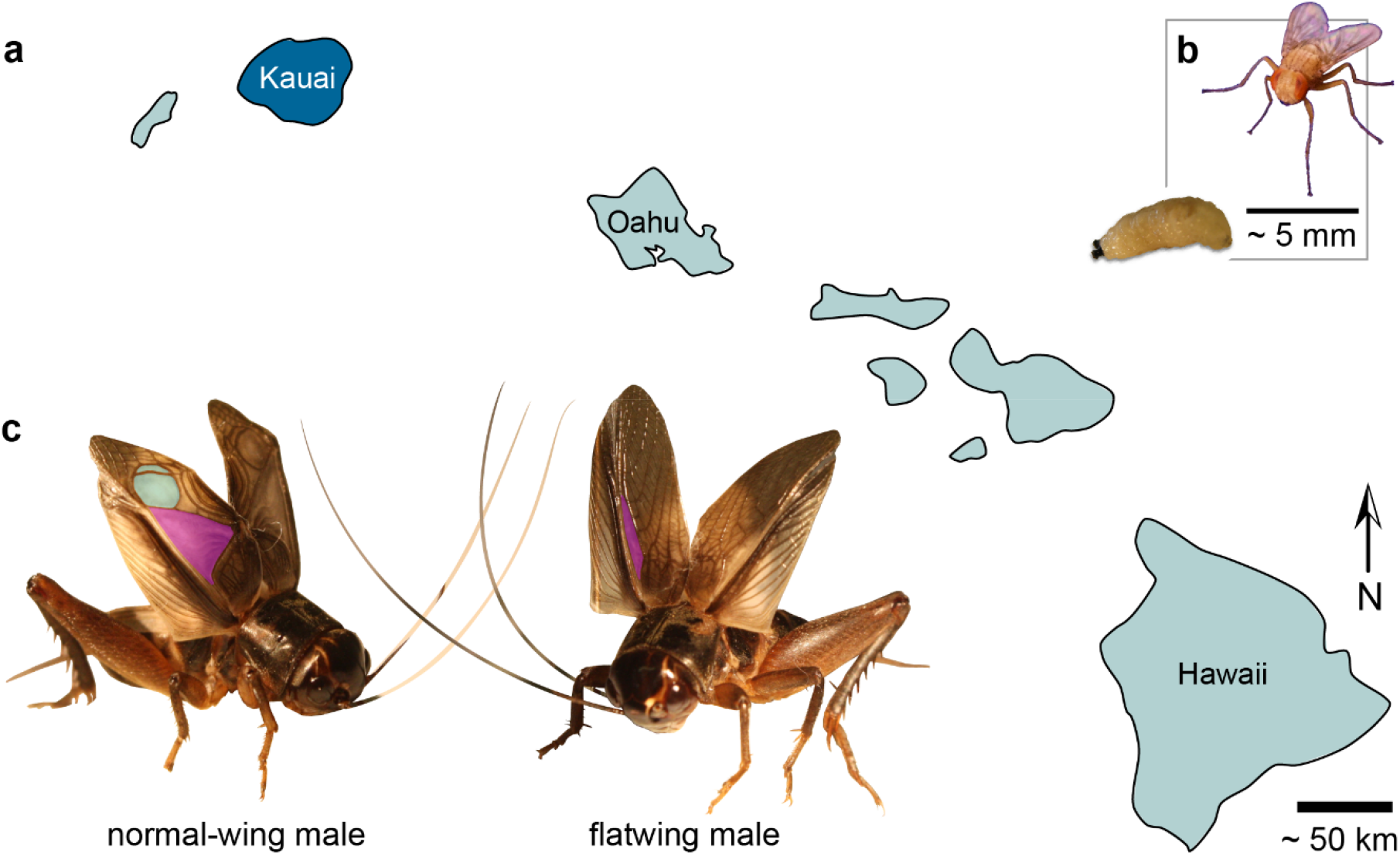
Rapid evolutionary loss of song in Hawaiian crickets. **a,** The field cricket *T. oceanicus* is thought to have migrated to the Hawaiian archipelago from other islands in Oceania, and is attacked by the fatal, acoustically-orienting parasitoid fly *Ormia ochracea* on Kauai, Oahu and Hawaii. We studied crickets from a population in Kauai, highlighted in dark blue, where parasitoid infestation rates have historically been highest. **b**, Adult female fly and mature parasitoid larva. Gravid female flies locate hosts by eavesdropping on singing male crickets, then they eject larvae that burrow into the host and consume its viscera before emerging to pupate. Infestation is fatal, and the flies exert significant natural selection against male song. **c**, Normal-wing males (left) of this field cricket species produce advertisement, courtship and aggressive songs by elevating and rubbing together forewings that bear specialised sound-producing venation. A toothed file on the right wing engages with a thickened ridge of tissue on the opposite, causing resonators to vibrate and produce sound. Two principal resonators are highlighted on this male’s right forewing: the harp in purple and the mirror in turquoise. Flatwing males (right) have wings that are feminised and lack, or have severely reduced, resonators. They still make wing motions characteristic of singing despite the structural inability to produce sound^93^, but their silence protects them from the fly^4^. The flatwing phenotype segregates as a single-locus mutation on the X chromosome, and 100% of males from the population studied on Kauai now exhibit flatwing morphology. (Photo credits: N.W. Bailey)

The locus controlling the expression of flatwing morphology could have arisen through *de novo* mutation(s) coinciding with the time of the phenotype’s first observation in 2003, it could have invaded the genome of the Kauai population via migration from an unknown location elsewhere in Hawaii (flatwing morphs have not been observed outside of the Hawaiian islands), or it could have existed for much longer in the population but at extremely low levels, evading detection by researchers. Studies of insecticide resistance in insects and of melanic morphs of Lepidoptera provide some precedent. While some museum specimens collected before the invention of organophosphates have been shown to contain insecticide-resistance alleles^10^, in other cases, resistance alleles arose *de novo*, and also invaded populations and spread under selection^11^. In the peppered moth, a canonical example of rapid evolution in the wild, melanism had a single recent origin approximately corresponding to the start of the industrial revolution^12,13^, but melanic morphs are common in many insects and may persist at low frequencies due to negative pleiotropy, at least until favourable selective conditions occur^14^. In *T. oceanicus*, parasitoid pressure pre-dated the appearance of flatwing in the Kauai population^4^, thus the *de novo* or introduction scenarios are most plausible.

We studied the genomic signature of song loss in the population on Kauai where flatwing crickets were first discovered, and in which rapid spread has been most thoroughly documented^4^. We sequenced the *T. oceanicus* genome, generating an assembly of 2.045 Gb consistent with flow cytometry size estimates^7^, with a scaffold N50 of 62.6 kb (Extended Data Table 1). We established an F_3_ mapping population using crosses designed to maximise recombination on the X chromosome, which is only diploid in females (Extended Data Fig. 1). Mapping offspring and parents were sequenced using RADseq, and a map was assembled containing 19 linkage groups. *T. oceanicus* has a haploid chromosome number of (13+X). We identified linkage group 1 (LG1) as the X chromosome by applying coverage and heterozygosity filters and dummy coding putative X-markers prior to constructing the map. LG1 was the largest linkage group, with a female recombination length of 379 cM and a male length of 195 cM (Extended Data Fig. 2). After resolving chimeric scaffolds (Extended Data Table 2), 35.6% of the genome was anchored to a linkage map using a LOD5 cutoff (Extended Data Table 3) (Fig. 2a).

**Fig. 2.**
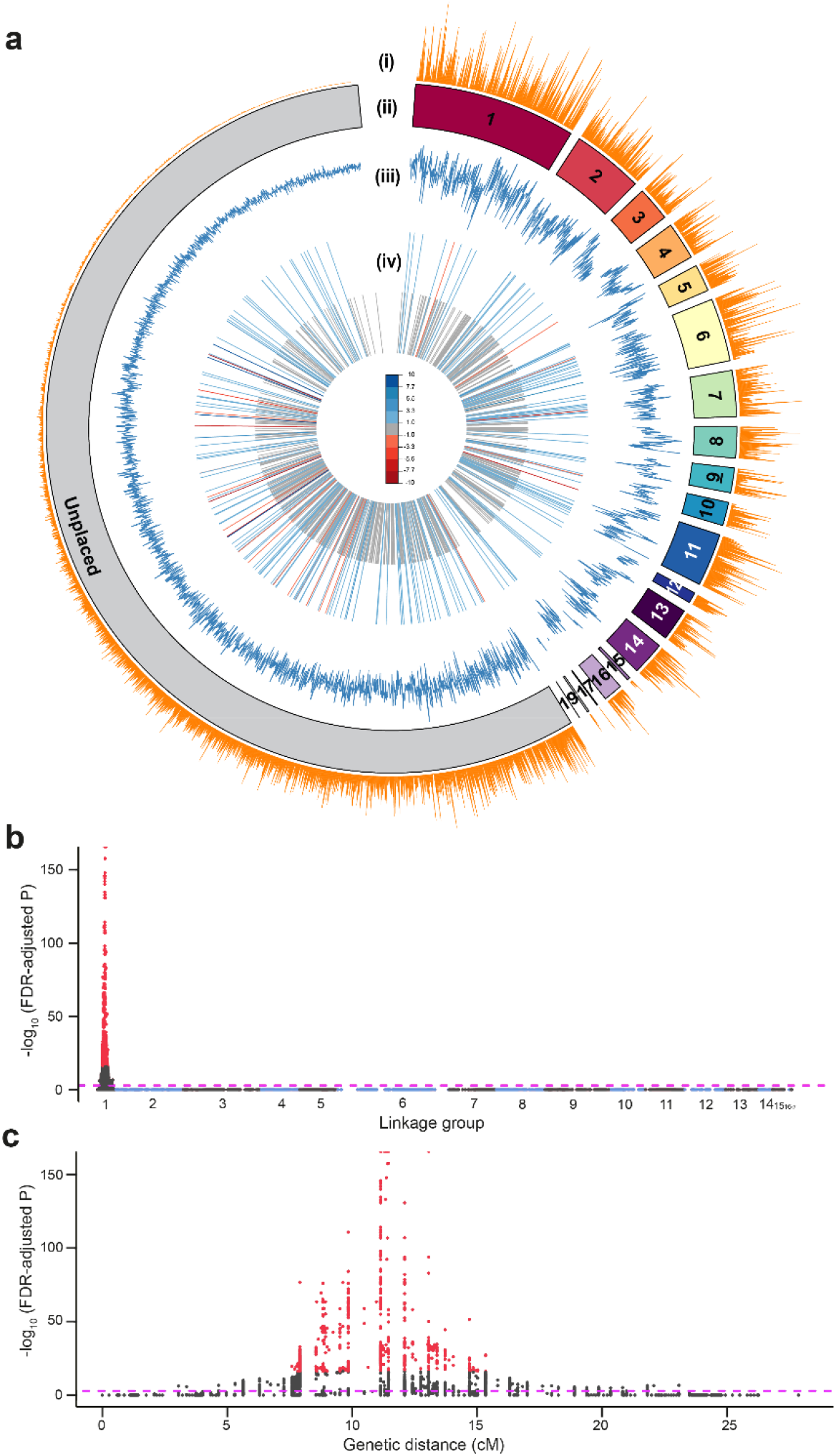
*Teleogryllus oceanicus* genome and regions associated with the flatwing phenotype. **a,** Circos plot providing an overview of the genome. Linkage groups (LGs) upon which genome scaffolds were anchored are shown in different colours, with unplaced scaffolds in gray. LG1 was identified as the X chromosome based on heterozygosity and coverage filters (see Main Text). Tracks (i): gene density, (ii): linkage group pseudomolecules, (iii): transposable element density, (iv): genes DE in the thoracic tissues of embryos homozygous for *flatwing* vs. *normal-wing* genotypes. Longer bars are DE genes for which log_2_FC > 1 between genotypes, and short grey bars are all other DE genes. Colours indicate the magnitude of upregulation (red) versus downregulation (blue) in *flatwing* compared to *normal-wing* embryos. **b,** Genome-wide Manhattan plot of the flatwing QTL. Alternating shades of grey and blue indicate different LGs. The horizontal dashed line indicates an FDR-corrected significance threshold of (*P* < 0.001), and the top 1% most significant QTL markers are plotted in red. **c,** Enlarged plot for LG1 (X chromosome) showing the flatwing-associated peak.

We performed gene prediction and annotation using custom pipelines incorporating *ab initio*, homology, and transcriptome-based approaches (Extended Data Fig. 3). Evidence from different gene prediction and annotation methods was weighted and filtered to predict a final, conservative set of 19,157 genes, 75% of which had functional annotation (Extended Data Table 4, Extended Data Fig. 4). Gene density was assessed (Fig. 2a track i), and we tested whether the putative X linkage group showed a different distribution of repeat content relative to the other linkage groups, across eight common categories of repeats. It did not (Fig. 2a track iii, Extended Data Table 5, Extended Data Fig. 5). *T. oceanicus* gene features were compared to 10 other insect species (Extended Data Table 6), and we contrasted transposable element classifications with three other recently published insect genomes (Extended Data Table 7). The *T. oceanicus* genome and metadata associated with it are curated in ChirpBase (www.chirpbase.org), a GenomeHubs Ensembl genome browser^15^ that we created as an openly available, community-based genomics resource for researchers working on singing insects.

*Flatwing* was definitively mapped to the putative X chromosome (Fig. 2b) using markers supported by a LOD10 cutoff and a mixed model, ANOVA-based approach designed to control for uneven genomic relatedness caused by family structure in the mapping crosses. To cope with the particularly high marker association on the putative X chromosome caused by the Mendelian mode of inheritance of *flatwing* and the different effective population size of the X compared to autosomes, we identified the QTL using only the top 1% of markers after FDR correction, yielding a prominent peak occupying approximately one third of the X chromosome (Fig. 2c). Flatwing morphology is observable in males during mid-to late-instar stages of juvenile development, so we examined early embryonic gene expression differences associated with *flatwing*. Females carrying the genotype cannot be visually distinguished and embryos cannot be readily sexed, so we used replicate laboratory lines homozygous for *flatwing* or *normal-wing* genotypes to detect widespread differential gene expression in the developing thoraces of embryonic crickets. We found 830 genes differentially expressed (DE), 204 of which had a log_2_ fold-change > 1, and a predominant pattern of down-regulation in *flatwing* crickets (Extended Data Table 8, Extended Data Fig. 6). DE genes associated with *flatwing* were widely distributed across linkage groups and unmapped scaffolds (Fig. 2a track iv).

These physically dispersed expression effects suggest that *flatwing* acts as a master regulatory switch during early development, with a broad cascade of downstream effects. Pathways reconstructed using differential expression data are consistent with such a mode of action. For example adherin junction activity was enriched, which affects epithelial patterning during early development (Extended Data Tables 9 & 10). Using a stringent and redundant approach combining information from gene sets identified in the QTL study, RNA-seq experiment and a previously-published bulked segregant analysis^7^, we identified 51 annotated protein-coding genes located within LG1 as top flatwing-associated candidates (Extended Data Table 11). GO enrichment analysis indicated that *positive regulation of developmental process* was overrepresented in this candidate gene set, with three genes in particular (*NBL1, GOGA4, UNC89*) known to play a fundamental role in the regulation of cell differentiation (Extended Data Table 12).

In most pterygote insects, wings are derived from imaginal discs formed during development by the invagination of embryonic ectoderm^16^. Previous work mainly in *Drosophila melanogaster* has established that the developmental elaboration of wing venation patterns requires the involvement of numerous transcription factors and complex coordination across numerous signalling pathways^17^. Here, we found that 8 of 51 flatwing associated candidate genes have reported involvement in *D. melanogaster* wing development. For example, *stat92E* expands the proximodistal axis of the wing imaginal disc, subdividing and patterning it^18^. *Collier* encodes a transcription factor required for wing disc patterning^19^, and *Myoglianin* expression is required for normal wing disc development^20^. *ROR1* encodes a transmembrane tyrosine-protein kinase receptor involved in phosphorylating MAP kinases^21^, and reduction of MAPK activity through *ROR1* silencing can lead to a loss of wing venation phenotype^17^. The protein krasavietz is encoded by *PKRA*, and establishes planar cell polarity in the wing^22^, disruption of which can lead to wing distortion^23^. Knockouts and mutants in *Pelle*, *Gen5*, and *Plexin-A4* show wing shape and venation alterations with features similar to flatwing^24-26^.

We tested the consequences of the rapid invasion of *flatwing* into the *T. oceanicus* genome by focusing on a distinct, close-range sexual signalling modality that operates alongside acoustic signalling in field crickets. Cuticular hydrocarbons (CHCs) are long-chain, waxy molecules expressed on insect cuticles. CHCs are thought to have evolved for dessication resistance, and they tend to be expressed as a bouquet of numerous individual hydrocarbon compounds. *T. oceanicus* CHCs are sexually dimorphic and function as sexual signals during male and female mate choice^27-29^, and they have been found to vary between flatwing and normal-wing male crickets^30^. We characterised the CHC profiles of F_3_ mapping individuals, all of which were raised in a common garden environment, by extracting their CHCs and using gas chromatography – mass spectrometry (GCMS) to measure the abundance of 26 individual compounds (Fig. 3a) (Extended Data Table 13). By performing dimension reduction using principal components (PC) analysis of the CHC profiles, we first established that, in our mapping population, males carrying *flatwing* showed noticeable and significantly different CHC profiles from *normal-wing* males (Fig. 3b) (multivariate analysis of variance on 6 principal components with eigenvalues > 1 describing male CHC blends: F_6,191_ = 29.769, p < 0.001) (Extended Data Table 14).

**Fig. 3.**
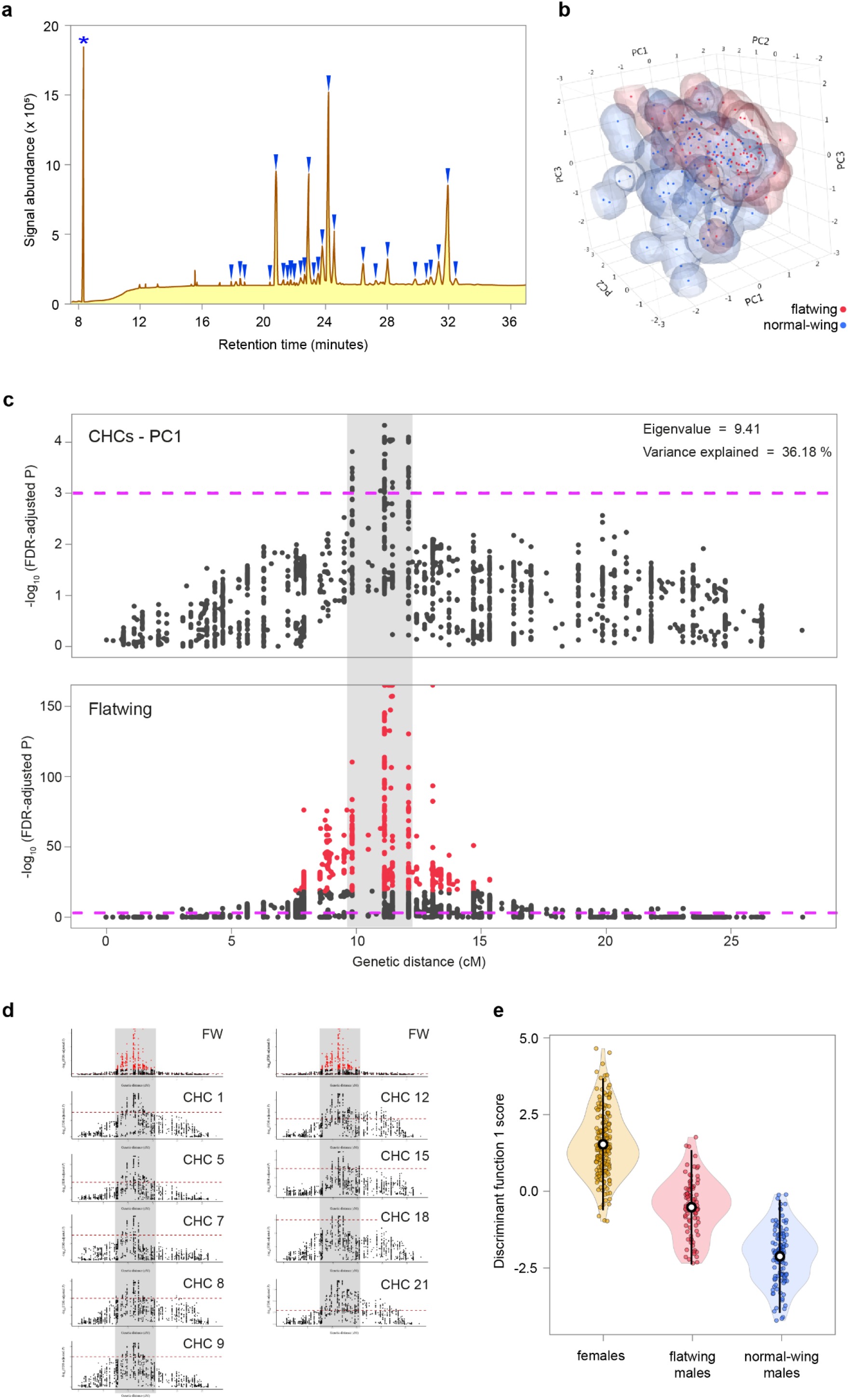
Pleiotropic effects of *flatwing* cause feminisation of chemical sexual signals. **a,** Diagram of a *T. oceanicus* cuticular hydrocarbon (CHC) chromatogram, with the 26 measured peaks indicated by blue wedges. The asterisk indicates the internal standard (pentadecane). **b,** Space-filling scatterplot of the first three principal components describing male CHC profiles, illustrating differences between flatwing and normal-wing males (variance explained for PC1: 35.18%, PC2: 10.14%, PC3: 9.58%). **c,** Comparison of QTL on the putative X chromosome for CHCs (top; first principal component mapped) and flatwing (bottom, same as Fig. 2C). Grey shading indicates the extent (in cM) of the CHC peak, showing overlap with the flatwing QTL. Dashed lines indicate FDR-corrected significance of p < 0.001, red points the top 1% significant flatwing QTL markers. Note the different y-axis scales. **d,** Univariate analyses revealed nine individual CHC components which also co-localised with flatwing. The original flatwing QTL is plotted at the top of each column. Grey shading spans the genetic region of co-localisation. Numbers refer to compounds indicated in **a**, and dashed lines indicate an FDR-corrected significance threshold of p < 0.001. **e,** Discriminant function scores describing variation in CHC profiles among female, flatwing male and normal-wing male mapping individuals. Discriminant function 1 explained 78.8% of the variance in CHC profiles between groups. Means ± 2 s.d. are indicated by open black-and-white circles and lines, respectively.

QTL analysis was then performed on the first six CHC PCs to determine whether *flatwing*-associated variation in male CHC profiles mapped to identifiable genomic regions. The putative X chromosome, LG1, was of particular interest, because we hypothesized that the striking variation between CHC profiles of flatwing and normal-wing males could be a pleiotropic effect of *flatwing*. Genetic mapping of CHCs was performed blind to male morphotype. PC1, which explained over a third of the variance in male CHC profiles, mapped to a ca. 2.5 cM region strongly co-localised with *flatwing* (Fig. 3c). PCs 4 and 6 also had co-localizing peaks (Extended Data Fig. 7). As dimension reduction for CHCs can obscure phenotypic patterns in the original individual chemical compounds, we mapped each of the 26 compounds separately. Of these, 9 showed significant peaks co-localising with *flatwing* (Fig. 3d). We recovered no autosomal QTL peaks for PCs 1-6, and only one QTL peak for one compound on one autosome (compound 11, 7-C31ene, on LG8). However, the latter peak was weakly supported, with only a single marker showing an association at FDR-corrected p < 0.001.

We interrogated genes on scaffolds under the CHC QTL peaks following a similar procedure used to produce the *flatwing* candidate gene set (Extended Data Table 15). Of 55 protein-coding genes, a subset of 6 were implicated for every CHC trait with a significant QTL peak, and these 6 genes were also present in the *flatwing* candidate gene set. These are strong candidates for testing the pleiotropic consequence of evolved acoustic sexual signal loss on chemical sexual signals. Our final step was to explore the nature of the phenotypic shift in flatwing male CHC profiles. It is unknown how flatwing males’ profiles compare to those of females^30^, but given the generally feminising effect of *flatwing* on male wing morphology, we predicted that flatwing males’ CHC profiles would also be feminised. We compared them to the profiles of normal-wing males and females using discriminant function analysis on profiles from all three groups. Discriminant function 1 (eigenvalue = 2.526) explained 78.8 % of the variance, and indicated that flatwing male crickets’ CHC profiles are strongly feminised (Fig. 3e). Their CHCs appear to be correspondingly less attractive to females^31^.

The rapid emergence and spread of flatwing crickets on Kauai has been described as one of the fastest rates of evolutionary adaptation ever documented in the wild^32^. Nearly a century ago in 1930, R. A. Fisher^8^ developed a ‘geometric’ model that describes the genomic landscape of such early-stage adaptation and predicts what mutational features are associated with adaptive change. In doing so, he reconciled the prevailing, gradualist view of evolution with seemingly inconsistent units of discrete Mendelian inheritance that were being discovered and characterised at the time. Fisher’s key insight was that the process of evolutionary adaptation tends to favour mutations of small effect, with impacts narrowly limited to the phenotypic variants directly under selection^33^. However, he built exceptions to this general rule into his model when selection is severe, and the genomic signature of song loss in Hawaiian *T. oceanicus* uniquely confirms and illustrates this insight. Adaptation was recent, abrupt and proceeded rapidly in this system. Prior work on *T. oceanicus* has found differences in the level of phenotypic plasticity, gene expression, and other reproductive characteristics such as male testis size between male *normal-wing* and *flatwing* genotypes^34-36^, and our present findings reveal the genomic footprint of strong, associated effects on sexual signalling in an entirely different sensory channel. These consequences of rapid adaptive trait loss are early-acting, genome-wide, and impact a range of important fitness traits. The suite of characters affected in flatwing crickets is reminiscent of feminised alternative male morphs in ruff (*Calidris pugnax*) in which supergene architecture controls size, ornament and behavioural traits simultaneously^37^, and in feminised bulb mites^38^. What is surprising is that an evolved loss of function could lead to such similarly wide-ranging phenotypic impacts so quickly. The genomic signature of recent, rapid trait loss in *T. oceanicus* confirms Fisher’s predictions about adaptive evolution – by demonstrating the exception to his rule.

## METHODS

### Cricket rearing and maintenance

Laboratory stocks of *Teleogryllus oceanicus* were established from eggs laid by wild-caught females from a population on the Hawaiian island of Kauai in 2012, and a population near Daintree, Australia in 2011. Stocks were maintained in the laboratory within 16 L plastic containers containing cardboard egg cartons for shelter. All crickets were reared in a single, temperature-controlled chamber a 25 °C, on a 12:12 light:dark cycle. They were maintained regularly and fed *ad libitum* with Burgess Excel Junior and Dwarf rabbit pellets and provided water in a moist cotton pad that also served as an oviposition substrate. Throughout all experiments, all crickets were reared in a common-garden environment in the same temperature-controlled chamber.

### Genome sequencing

Three Illumina sequencing libraries were prepared using genomic DNA extracted from the head capsule and muscle tissue of a single *T. oceanicus* female using a DNeasy Blood & Tissue kit (Qiagen). The female was sourced from the Kauai stock population. gDNA was quality-checked using Nanodrop and Qubit prior to Illumina library preparation and sequencing at Edinburgh Genomics. We prepared three standard paired-end TruSeq libraries with insert sizes of 180 bp, 300 bp, and 600 bp. We supplemented reads from the above three Illumina libraries with additional sequences from two TruSeq Nano Pippin selected libraries with insert sizes of 350 bp and 550 bp, one 8 kb Nextera gel-plus mate-pair library, and 1 PacBio library. For these libraries, gDNA from a separate, single female cricket from the same laboratory population was extracted using a high molecular weight Genera Puregene Cell Kit (Qiagen). The first three TruSeq libraries were sequenced on 5 lanes of an Illumina HiSeq 2000 v3 to yield 100 bp paired-end reads. NanoPippin libraries and the Nextera mate-pair library were sequenced on 2 Illumina HiSeq 2500 lanes to yield 250 bp paired-end reads. To construct the PacBio library, we purified the extraction with 1x AMPure beads (Agencourt) and performed quality control using Nanodrop and Qubit. Average DNA size and degradation was assessed using a high sensitivity genomic kit on a fragment analyzer. Size-selected and non-size-selected libraries were made by shearing gDNA using g-TUBEs (Covaris). Size selection was performed using the BluePippin DNA Size Selection System with 0.75% cassettes and cutoffs between 7 and 20 kb. Preparation of both libraries then proceeded using the same protocol. We treated DNA for 15 min at 37 °C with Exonuclease V11. DNA ends were repaired by incubating for 20 min at 37 °C with Pacific Biosciences damage repair mix. Samples were then incubated with end repair mix for 5 min at 25 °C followed by washing with 0.5x AMPure and 70% ethanol. DNA adapters were ligated overnight at 25 °C. Incubation at 65 °C for 10 min was used to terminate ligation reactions, and then samples were treated with exonuclease for 1 hr at 37 °C. We purified the SMRTbell library using 0.5x AMPure beads and checked quality and quantity using Qubit assays. Average fragment size was quantified using a fragment analyser. For sequencing, primers were annealed to the SMRTbell library at values determined using PacBio’s Binding Calculator. A complex was formed using DNA polymerase (P6/C4 chemistry), bound to MagBeads, and then used to set up 43 SMRT cells for sequencing to achieve 10X coverage. Sequencing was performed using 240 min movie times.

### Genome assembly

Raw reads from all Illumina libraries were trimmed using cutadapt v.1.8.3^39^ to remove adapters, primers and poor quality bases, and then error-corrected using BLESS^40^. PacBio reads <1,000 bp were discarded. The original fragment length of the 350 bp library was shorter than the sequenced paired read length of 500bp, so reads from this library were merged using Vsearch v.1.10.1^41^. Platanus v.1.2.4^42^ was used to assemble error-corrected reads from all Illumina libraries except the mate-pair library; reads from the latter were added at the scaffolding stage. Next, we selected the contigs >1,000 bp and combined these with the PacBio data to generate a hybrid assembly using PBJelly v.15.2.20^43^. Pilon v.2.1^44^ was used to improve local base accuracy, and BUSCO v.2.1^45^ was used to assess genome quality through gene completeness.

### Repeat annotation

We used *de novo* and homology-based approaches to identify repetitive regions. We first built a *de novo* repeat library using RepeatModeler^46^, with dependencies RECON and RepeatScout^47^. To scan and classify interspersed repeats and low complexity DNA sequences at the DNA level, we searched the cricket genome sequence against the Dfam consensus database^48^, RepBase^49^, and the *de novo* repeat library using RMBlast^50^ and RepeatMasker^51^. Protein-level repeats were identified by searching against the TE Protein Database using RepeatProteinMask^51^.

Unclassified repetitive elements were further classified by TEclass^52^, a programme using a support vector machine learning algorithm. Tandem repeats were also identified in the cricket genome using Tandem Repeat Finder^53^.

### Gene prediction

Before running gene prediction pipelines, repetitive regions identified above were masked using an in-house Perl script. We built a pipeline including *ab initio*, homology and transcriptome-based methods to predict protein-coding genes in the cricket genome (Extended Data Fig. 3). For *ab initio* prediction, SNAP^54^, Glimmer-HMM^55^, GENEID^56^, and BRAKER1^57^ were used to generate preliminary gene sets from the repeat-masked genome. Specifically, reads obtained from the *T. oceanicus* transcriptome were aligned against the repeat masked genome with TopHat2^58^. SAMTOOLS^59^ was used to sort and index the resulting Binary Alignment Map (BAM) format file.

This BAM file was processed in BRAKER1^57^, which used transcriptome data to train GENEMARK-ET^60^, generate initial gene structures, and then subsequently train AUGUSTUS^61^ and finally integrate RNA-seq information into final gene predictions. For other *ab initio* gene prediction programmes, gene sets from *Locusta migratoria*^62^, *Acyrthosipon pisum*^63^, and *Drosophila melanogaster*^64^ were used for model training. For homology-based prediction, we aligned protein sequences of five insect species (*L. migratoria*^62^, *Drosophila melanogaster, Anoplophora glabripennis*^65^, *Nilaparvata lugens*^66^, and *Cimex lectularius*^67^) to the repeat-masked cricket genome using TBLASTN (E < 10^-5^)^50^. The boundaries of potential genes were further identified using BLAST2GENE^68^. We then ran GENEWISE2^69^ to obtain accurate spliced alignments and generate a final, homology-based gene set. For prediction based on transcriptome data, the *de novo* transcriptome assembly generated by Trinity^70^ was filtered based on gene expression level, and then passed to Program to Assemble Spliced Alignments (PASA)^71^. PASA performed transcript alignments to the cricket genome, generated a new transcript assembly, and predicted gene structures. All gene sets predicted by *ab initio*, homology, and transcriptome-based methods were then combined into a weighted consensus gene set using EVidenceModeler (EVM)^72^. We removed genes likely to be spurious, those with low EVM support, partial genes with coding lengths shorter than 150 bp, and genes only supported by a minority (≤ 2) of *ab initio* methods^73^. PASA was used to further update the filtered consensus gene set to produce a finalised official gene set. The completeness of this final gene set was assessed by both BUSCO v.2.1 (using the arthropoda dataset)^45^ and transcriptome data.

### Functional assignment

Putative gene functions were assigned based on InterPro^74^, SwissProt^75^, TrEMBL^75^ and RefSeq non-redundant (NR) protein and Kyoto Encyclopedia of Genes and Genomes (KEGG) gene databases. Briefly, we first obtained protein sequences from the final gene set using EVM^72^. Functional annotation and gene ontology terms were assigned to genes based on protein sequence, using InterProScan 5^76^. These proteins were also blasted against SwissProt, TrEMBL and NR databases (PLASTP, E < 10^-5^), and assigned their best hits as functional annotations. Gene ontology (GO) terms were assigned using GO annotations downloaded from the GO Consortium^77,78^. BLAST2GO^79^ was implemented to further assign unassigned genes using NCBI databases, and KEGG Orthology (KO) terms were assigned using BlastKOALA^80^.

### Genome anchoring

ALLMAPS^81^ was used to detect chimeric scaffolds, anchor the cricket genome to the linkage map (see below), and construct pseudo-molecules (reconstructed portions of chromosomal sequence). We first built a consensus genetic map based on male and female genetic distances obtained from linkage maps, in which equal weighting was applied for both sexes. Then, scaffolds for which more than four markers mapped to multiple linkage groups were designated as chimeric scaffolds, and split. After this correction was applied, scaffolds anchored to the linkage maps were oriented and ordered based on the consensus genetic map. We used a custom Perl script to order unanchored scaffolds according to their length, and liftOver^82^ to convert genome coordinates based on anchoring results.

### Genome browser development (ChirpBase)

We created ChirpBase, an open-access community genomics resource for singing insects, such as field crickets and katydids. The resource can be accessed at www.chirpbase.org where users may view and download genomic data and scripts presented in this study in addition to uploading data. An index page links to an ensembl page, where assembly statistics can be visualised using a Challis plot or compared in tabular format. A plot illustrating codon usage is presented, as well as a visualisation of BUSCO scores. Additional pages linking from this include a basic local alignment search tool (BLAST) page and a download page where raw data can be accessed. We used the GenomeHubs framework to set up ChirpBase^14^. Briefly, the databased is hosted using a Linux container (LXC) on a remote computer, linked to a cluster via an intermediate import computer. A MySQL docker container was started in the LXC, where a database *ini* file resided to guide additions to the database. An Ensembl-easy mirror Docker container was run to import the database into the MySQL container, uploading data designated in the *ini* file from the LXC to the database. The MySQL container links to an Ensembl EasyMirror container, BLAST container, and a download container.

### Linkage and QTL mapping crosses

We constructed a linkage map for *T. oceanicus* using a series of crosses to maximise recombination on the X chromosome (Extended Data Fig. 1), combined with restriction-site associated DNA sequencing (RAD-seq) to identify markers. *Flatwing* segregates on the X chromosome in both Kauai and Oahu populations^6,7^, so mapping was performed with F_3_ offspring to increase recombination on the X. We set up two parental mapping families by crossing a flatwing sire from the Kauai population with a virgin dam from the Daintree, Australia population. Daintree females were used in the cross because flatwings do not exist in that population, and other sexually-selected traits such as song and cuticular hydrocarbons show significant divergence between Australian and Hawaiian populations^83^, which maximised our opportunity to genetically map segregating variation in other phenotypes. Female F_1_ offspring from parental crosses were heterozygous for *flatwing*, enabling recombination on the X. Full-sib matings were then performed with F_1_ males, all of which were normal-wing. The resulting F_2_ female offspring were a segregating mix of homozygous *normal-wing* genotypes on the X, or heterozygous with respect to wing morph. Recombination between *flatwing* and *normal-wing* genotypes was similarly possible in the heterozygous F_2_ females, but their phenotype is not externally detectable. To further increase recombination on the X, we performed another generation of crossing by mating F_2_ females with full-sib *flatwing* males from the same generation. Screening male morph types in the resulting F_3_ offspring enabled us to identify F_2_ crosses involving heterozygous females, as all male offspring of homozygous *normal-wing* females expressed normal-wing morphology. The crossing procedure resulted in 10 F_3_ mapping families from the original two parental families, for a total of 192 females, 113 normal-wing males, and 86 flatwing males.

### Marker identification using RAD-seq

RAD-seq was used to identify single nucleotide polymorphisms (SNPs) in F_3_ offspring (n = 391), P_0_ dams and sires (n = 4), and the F_2_ sires and dams (n = 19) that were used to produce mapping individuals in the F_3_ generation. For each individual, gDNA extraction and quality control was performed as described above prior to library preparation. gDNA was digested using SbfI (New England BioLabs). We barcoded individuals by ligating P1 adapters (8 nM), then sheared and size selected 300-700 bp fragments. After ligating P2 adapters to sheared ends, parents were sequenced to an average coverage of 120x and offspring to 30x on an Illumina HiSeq 2000.

### Construction of linkage map

Reads from all paired end RAD libraries were demultiplexed by sample using process_radtags from Stacks^84^, mapped against the reference genome assembly using BWA-MEM^85^ and duplicates marked using PicardTools MarkDuplicates (http://broadinstitute.github.io/picard). Variants were called using samtools mpileup (version 1.3, parameters -d 2000 -t DP,DPR,DV,DP4,SP -Aef -gu) and bcftools call (version 1.3, parameters -vmO z -f GQ). The resulting variants were filtered using vcfutils.pl (included with bcftools) with minimum quality 50 and a minimum read depth of 150 (-Q 50 -d 150) to only retain high quality variants. The vcf format was converted to the required lepmap2 input format using a custom script of the RADmapper pipeline (RAD_vcf_to_lepmap_with_sexmarker_conversion.py, https://github.com/EdinburghGenomics/RADmapper). During this conversion samples that did not fit relatedness expectations and samples from family J (which lacked a genotyped father) and P0 parents were excluded from linkage map creation. Putative X-linked markers (male_het <=1, female_het > 20, het_sire <=1) were converted to biallelic markers in the relevant male offspring and sires using a dummy allele (Extended Data Table 17). The linkage map was then created using the following steps and parameters in lepmap2 (Filtering: dataTolerance 0.05 keepAlleles=1; SeparateChromosomes: losLimit=10 sizeLimit=10 informativeMask=3;JoinSingles: lodLimit=5;OrderMarkers: filterWindow=10 polishWindow=100; OrderMarkers evaluateOrder: filterWindow=10 polishWindow=100). The resulting linkage map files were merged with the marker and sample information using a custom script from the RADmapper pipeline (LG_to_marker.py).

### QTL mapping

To identify the flatwing locus on the putative X chromosome (LG1), we performed ANOVA for each marker using the lm package in R (v. 3.1). Individual p-values were corrected to account for multiple testing using Bonferroni correction and markers supported by a LOD10 cutoff were plotted. QTL for all 26 cuticular hydrocarbon (CHC) peaks as well as the principle components from the CHC analysis were mapped to the linkage groups using mixed linear models in ASReml 4. Mapping used a GWAS-type approach, taking into account genetic relatedness between individuals^86^.

The markers mapped to the autosomal linkage groups 2-19 were filtered to contain only bi-allelic SNP markers with a MAF <=0.01 and <5% missing samples per marker. In addition, all grandparental, parental and female samples were removed together with samples that clustered with the wrong family or did not have CHC data. Only male samples were selected, as our aim was to map male CHCs (not sex-related associations) on the putative X (LG1) and autosomes using principle components from the CHC analysis as well as individual compounds as traits. The remaining 21,047 markers were used to calculate pairwise genetic relatedness with the first normalisation^87^. The resulting inverse relatedness matrix was used as random effect in a model: CHC trait ~ mu marker r! Giv(animal). P-values for all markers were extracted from the results and corrected for multiple testing using Bonferroni correction. The same model was used to assess LG1 separately with the same set of samples, for which 6,537 markers were used after filtering.

### Pure-breeding lines and embryo sampling for RNA-seq

Lines homozygous for the *flatwing* and *normal-wing* genotypes were produced following previously described methods^34^. Briefly, one generation of crosses was performed, starting with the laboratory population derived from Kauai and crossing males of either wing phenotype to virgin females of unknown genotype. Because the phenotypic effects of *flatwing* are sex-limited, family-level screening of the resulting male offspring was performed to select homozygous *flatwing* and homozygous *normal-wing* lines, resulting in a final selection of three pure-breeding lines for each morph genotype. Developing embryos were sampled from eggs laid by females from each line. Females were maintained in laboratory culture as above, and their oviposition substrates were monitored. Eggs were removed from the substrate and immediately preserved in 500 μL of RNAlater (Qiagen) at the stage when eye pigmentation first develops, ca. 2 weeks after laying. This time point corresponds approximately to embryonic stage 13-14 in the related grylline species *Gryllus bimaculatus*^88^. After removing the outer egg chorion, the thoracic segment of each nymph was microdissected. Nymphs cannot be sexed based on external morphology until a later stage of juvenile development, and as chromosomal sex determination is XX/XO, screening for sex-specific markers is not possible. To minimise potential variation in sex ratio of samples between lines, and ensure a sufficient volume of tissue to extract RNA, thoracic tissue from n = 8 nymphs was pooled for each replicate, and 6 biological replicates were produced for each morph type (2 per line).

### RNA-seq and gene expression profiling

Total RNA was extracted using the TRIzol plus RNA purification kit (Life Technologies) and DNAse treated using PureLink (Invitrogen). RNA was quantified and quality checked using Qubit assessment (Invitrogen) and Bioanalyser RNA Nano Chips (Agilent), respectively. To isolate mRNA we depleted samples with RiboZero. After verifying depletion, cDNA libraries were constructed using the ScriptSeq protocol (Epicentre) with AMPure XP beads for purification. Following barcoding and multiplexing, final quality was checked and qPCR performed using Illumina’s Library Quantification Kit (Kapa). Sequencing was performed on an Illumina HiSeq 2000 v3, with 1% PhiX DNA spike-in controls to produce 100 base paired-end reads. CASAVA v.1.8.2 was used to demultiplex reads and produce raw fastq files, which were then processed with Cutadapt v.1.2.1^38^ and Sickle v.1.200^89^ to remove adaptor sequences and trim low-quality bases. Further quality assessment was performed in FastQC. Expression analysis of RNA-seq data was performed broadly following the protocol published by Pertea et al. (2016)^90^. Reads were aligned to the genome using HISAT2 with strand-specific settings, and transcripts compiled for each sample in StringTie, using the gene annotation file as a reference, which were then merged across all samples to produce a single annotated reference transcriptome. Sample transcript abundances were estimated with the parameter -e specified to restrict abundance estimation to annotated transcripts. Differential expression analysis was performed at the gene level following normalisation of counts by trimmed mean of M-values (TMM), using a generalised linear model (GLM) with negative binomial distribution and a single predictor variable of ‘morph’ in the edgeR package^91^ in R v.3.4.1. Only genes with an expression level greater than 1 count per million in at least 3 samples were included in the analysis. Significance-testing was performed using likelihood ratio tests, and genes were considered significantly differentially expressed between morph genotypes if FDR-adjusted P-values were below a threshold of 0.05.

### Screening for top candidate genes associated with flatwing

We adjusted *P*-values for significant marker associations in the flatwing QTL mapping procedure using Bonferroni correction with a cut-off of *P* < 0.001. Three sources of information were used to comprehensively and robustly detect a set of top candidate genes associated with the flatwing phenotype. We detected genes (i.e. any overlapping portion of a predicted gene sequence cf. Extended Data Table 6) located in 1 kb flanking regions of all significant QTL markers after FDR correction as above, and defined these as QTL-associated candidates. We then subsetted these genes to retain only those located in the 1 kb flanking regions of the most significant (top 1%) of all QTL markers, and defined these as Top 1%-associated candidates. We also obtained the flatwing-associated sequences from a previously published bulk segregant analysis (BSA) of Kauai flatwings^7^, and defined the BSA reference sequences containing flatwing-associated SNPs as flatwing-associated BSA sequences. We mapped these BSA sequences to the *T. oceanicus* reference genome using BWA-MEM with default parameters^85^. Coordinates of mapped sequences were extracted from the resulting BAM files using SAMTOOLS^59^ and custom Perl scripts, and we only retained those sequences that were anchored to LG1. Genes within 1 kb of these retained sequences were defined as BSA-associated candidates. Finally, we extracted differentially expressed genes from the embryonic thoracic transcriptome analysis above, and defined these as DEG-associated candidates. To ensure a reliable final candidate gene set for flatwing, we only retained genes supported by at least two of these four gene sets. We used KEGG pathway mapping (colour pathway) to reconstruct pathways and obtain reference pathway IDs^92^. To characterise significantly enriched GO terms and KEGG pathways in DEGs, we implemented the hypergeometric test in enrichment analyses. *P* values for each GO and KEGG map term were calculated and FDR-adjusted in R.

### Cuticular hydrocarbon extraction and gas chromatography-mass spectrometry

We extracted CHCs from F_3_ mapping individuals prior to extracting gDNA for RADseq. Extraction and analysis of CHCs followed previous methodology^83^, which is briefly described here. Subjects were flash-frozen for several minutes at −20 °C and then thawed. They were individually placed into 4 mL borosilicate glass vials (QMX Laboratories) and immersed for 5 minutes in 4 mL of HPLC-grade hexane (Fisher Scientific), then removed from the vials and stored for later processing. We evaporated a 100 μL aliquot of each sample overnight in a 300 μL autosampler vial (Fisher Scientific). CHC extracts were transported to the University of Exeter for gas chromatography mass spectrometry (GC/MS) using an Agilent 7890 GC linked to an Agilent 5975B MS. Extracts were reconstituted in 100 μL of hexane with a 10 ppm pentadecane internal standard, and 2 μL of this was injected into the GC/MS using a CTC PAL autosampler at 5 °C. The carrier gas was helium and we used DB-WAX columns with a 30 m x 0.25 mm internal diameter and 0.25 μm film. Injection was performed in split-less mode. The column profile was optimised for separation of the CHC extract^83^ to start at 50 °C for 1 minute, followed by a temperature ramp of 20 °C per minute until finally holding at 250 °C for a total run time of 90 minutes. The inlet temperature was 250 °C and the MS transfer line was 230 °C. We recorded electron-impact mass spectra using a 70 eV ionization voltage at 230 °C, and a C_7_-C_40_ alkane standard was run as a standard to enable the later calculation of peak retention indices.

### Quantification and analysis of CHC profiles

For each individual, we used MSD CHEMSTATION software (v.E.02.00.493) to integrate the area under each of 26 CHC peaks (Extended Data Table 13) following Pascoal et al. (2016)^83^. Peak abundances were standardized using the internal pentadecane standard and log_10_ transformed prior to analysis. After accounting for samples that failed during extraction or during the GC run (n = 9), plus one normal-wing male CHC profile that was identified as an outlier and removed during analysis (Extended Data Fig. 8), we analyzed a total of n = 86 flatwing males, n = 112 normal-wing males, and n = 185 females of unknown genotype. To test whether CHC profiles differed between males of either wing morph, we first performed dimension reduction using principal components analysis (PCA) on male data only. JMP Trial 14.1.0 (SAS Institute Inc.) was used to draw a 3D scatterplot of the first three PCs. To assess statistical significance, we performed a MANOVA using all principal components with eigenvalue > 1.00 (n = 6). This indicated a highly significant difference among male morphs which formed the basis of QTL mapping described above. To visualise the difference between flatwing and normal-wing male CHC profiles with respect to female CHC profiles, we performed a discriminant function analysis (DFA) for all samples and all 26 peaks. DFA highlights the maximal difference between pre-defined groups, with maximum group differences indicated by the first DF axis. Statistical analyses of CHC data were done in SPSS (v.23).

### Data Availability

Raw reads from Illumina and PacBio genome sequencing libraries, embryo RNAseq reads, RADseq reads used in the linkage map and QTL analyses, CHC phenotype data will be made publicly available upon acceptance. Custom scripts are available online at http://www.chirpbase.org if not stated otherwise.

---

**Extended Data Table 1.** *T. oceanicus* genome metrics

| Maximum scaffold length (bp) | 2,637,780 |  |
| --- | --- | --- |
| Complete BUSCO (Ref total = 1,066) | 1,001 |  |
| % complete BUSCO (genome) | 93.9% |  |
| % complete BUSCO (gene set)<sup>a</sup> | 83.2% |  |
| Scaffold metrics | All contigs | Contigs > 1,000 bp |
| Total bases (gb) | 2,045,067,382 | 2,044,651,628 |
| N50 (bp) | 62,615 | 62,685 |
| Sequences in N50 | 6,139 | 6,136 |
| GC content (%) | 40 | 40 |
| Mean scaffold length (bp) | 10,335 | 10,355 |
| Sequencing library yields | Read pairs | Yield (GB) |
| Illumina TruSeq 180 | 277,076,641 | 55.42 |
| Illumina TruSeq 300 | 243,927,180 | 48.79 |
| Illumina TruSeq 600 | 238,275,727 | 47.66 |
| Illumina TruSeq Nano 350 | 70,959,741 | 14.19 |
| Illumina TruSeq Nano 550 | 63,415,263 | 12.68 |
| Illumina Nextera mate-pair | 229,431,023 | 45.89 |
| PacBio RSII | 5,771,779 | 21.74 |

<sup>a</sup> Final gene number identified in Extended Data Table 6

**Extended Data Figure 1.**
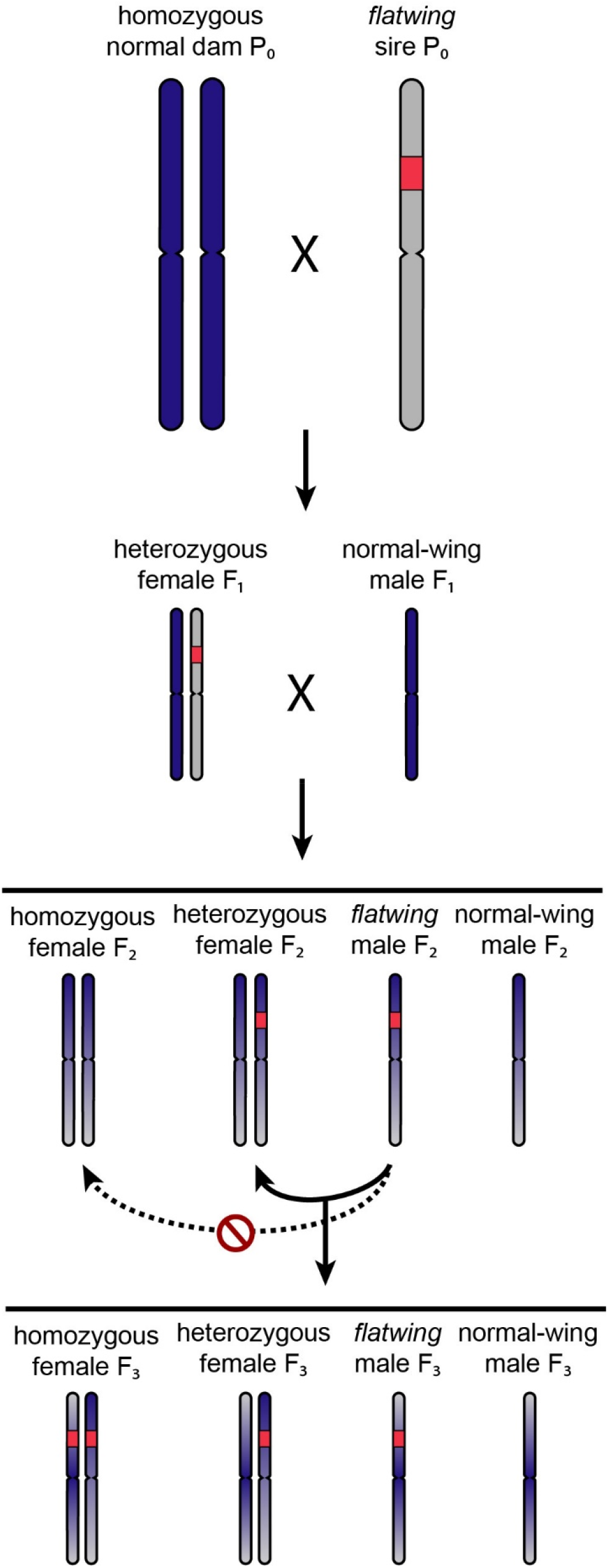
Cross design for linkage and QTL mapping. *Flatwing* segregates as a single-locus X-linked trait, so only X chromosomes are illustrated. A hypothetical *flatwing* locus is shaded in red. Females and males are XX/XO in *T. oceanicus*, so we performed three generations of crossing to enhance our ability to map *flatwing*. Homozygous *normal-wing* dams were obtained from a laboratory population of the same species originally derived from a population that has never contained *flatwing* (dark blue chromosomes). In the parental generation, these *normal-wing* dams were crossed to *flatwing* sires from Kauai (light grey chromosome, with hypothetical *flatwing* locus shaded in red). Dam genotypes were undetectable at generation F_2_ due to *flatwing’s* sex-limited expression, so only full-sib crosses for which the flatwing male phenotype segregated in the subsequent F_3_ generation were retained for phenotyping and QTL mapping (solid arrows).

**Extended Data Figure 2.**
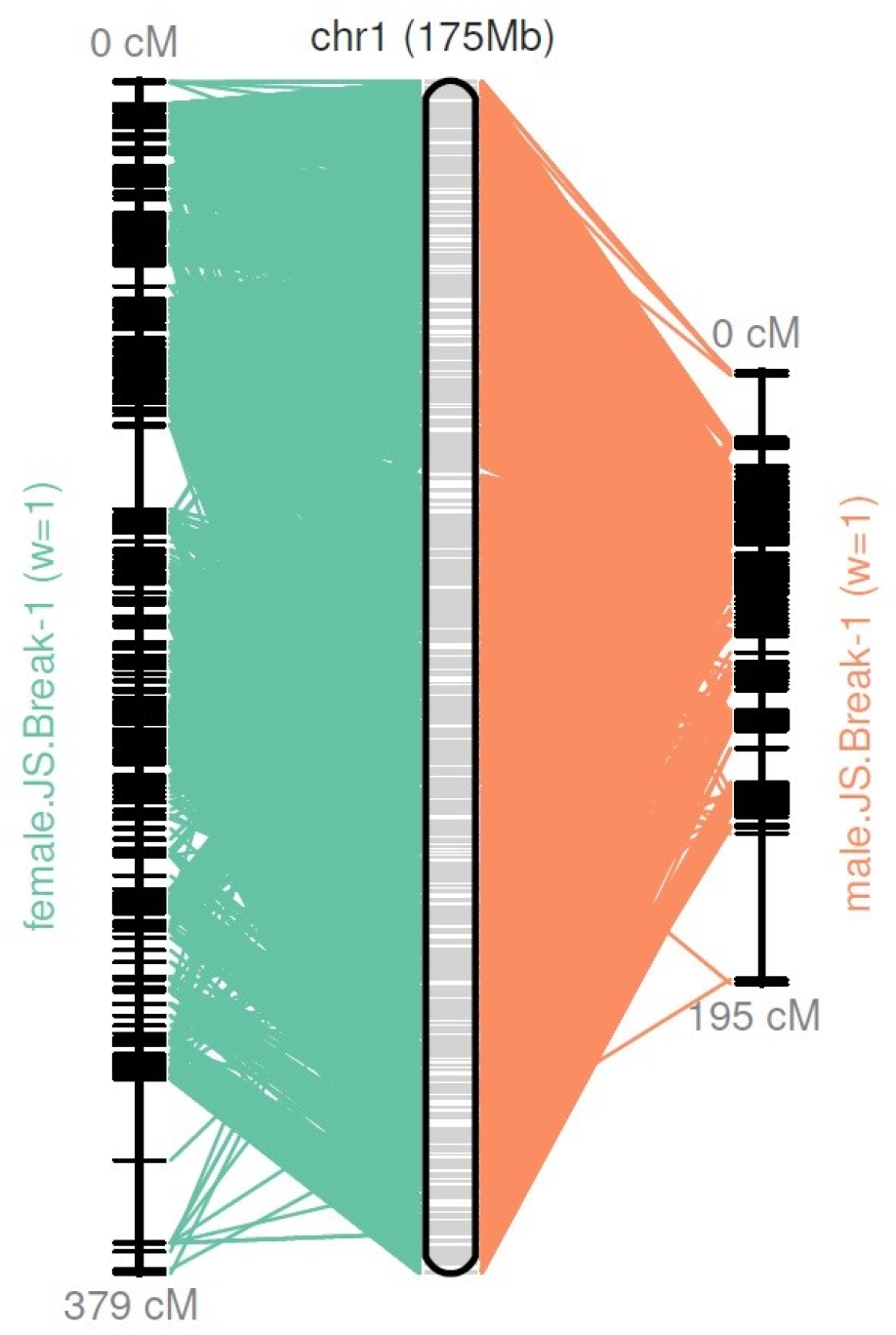
Reconstructed pseudomolecules for LG1 (putative X chromosome) using LOD5-supported markers. Female and male pseudomolecules are shown on the left and right, respectively, and lines connect the physical positions of markers on each pseudomolecule to map positions.

**Extended Data Table 2.**
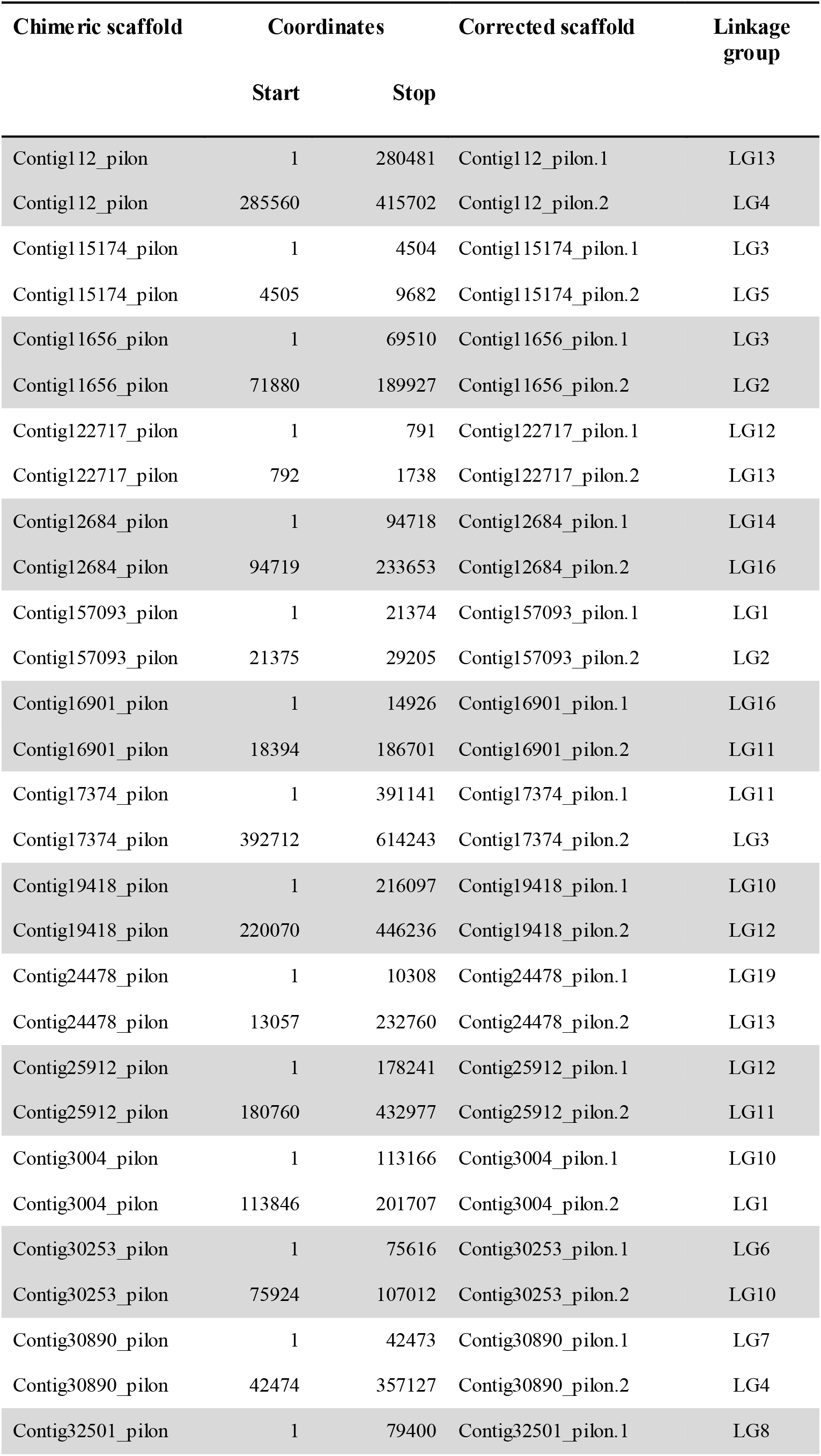
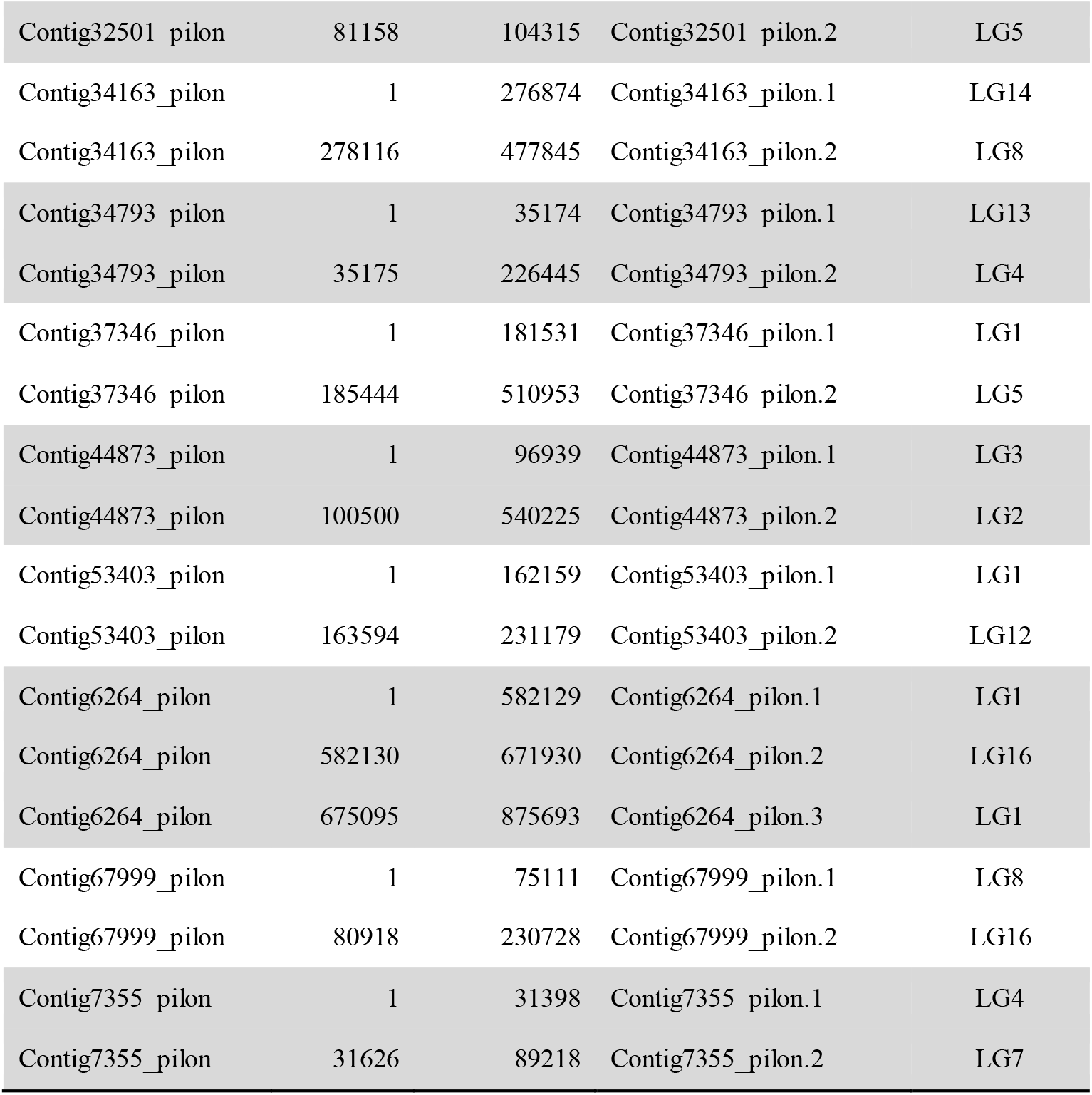
List of chimeric scaffolds identified and corrected in the *T. oceanicus* genome

| Chimeric scaffold | Coordinates |  | Corrected scaffold | Linkage group |
| --- | --- | --- | --- | --- |
|  | Start | Stop |  |  |
| Contig112_pilon | 1 | 280481 | Contig112_pilon.1 | LG13 |
| Contig112_pilon | 285560 | 415702 | Contig112_pilon.2 | LG4 |
| Contig115174_pilon | 1 | 4504 | Contig115174_pilon.1 | LG3 |
| Contig115174_pilon | 4505 | 9682 | Contig115174_pilon.2 | LG5 |
| Contig11656_pilon | 1 | 69510 | Contig11656_pilon.1 | LG3 |
| Contig11656_pilon | 71880 | 189927 | Contig11656_pilon.2 | LG2 |
| Contig122717_pilon | 1 | 791 | Contig122717_pilon.1 | LG12 |
| Contig122717_pilon | 792 | 1738 | Contig122717_pilon.2 | LG13 |
| Contig12684_pilon | 1 | 94718 | Contig12684_pilon.1 | LG14 |
| Contig12684_pilon | 94719 | 233653 | Contig12684_pilon.2 | LG16 |
| Contig157093_pilon | 1 | 21374 | Contig157093_pilon.1 | LG1 |
| Contig157093_pilon | 21375 | 29205 | Contig157093_pilon.2 | LG2 |
| Contig16901_pilon | 1 | 14926 | Contig16901_pilon.1 | LG16 |
| Contig16901_pilon | 18394 | 186701 | Contig16901_pilon.2 | LG11 |
| Contig17374_pilon | 1 | 391141 | Contig17374_pilon.1 | LG11 |
| Contig17374_pilon | 392712 | 614243 | Contig17374_pilon.2 | LG3 |
| Contig19418_pilon | 1 | 216097 | Contig19418_pilon.1 | LG10 |
| Contig19418_pilon | 220070 | 446236 | Contig19418_pilon.2 | LG12 |
| Contig24478_pilon | 1 | 10308 | Contig24478_pilon.1 | LG19 |
| Contig24478_pilon | 13057 | 232760 | Contig24478_pilon.2 | LG13 |
| Contig25912_pilon | 1 | 178241 | Contig25912_pilon.1 | LG12 |
| Contig25912_pilon | 180760 | 432977 | Contig25912_pilon.2 | LG11 |
| Contig3004_pilon | 1 | 113166 | Contig3004_pilon.1 | LG10 |
| Contig3004_pilon | 113846 | 201707 | Contig3004_pilon.2 | LG1 |
| Contig30253_pilon | 1 | 75616 | Contig30253_pilon.1 | LG6 |
| Contig30253_pilon | 75924 | 107012 | Contig30253_pilon.2 | LG10 |
| Contig30890_pilon | 1 | 42473 | Contig30890_pilon.1 | LG7 |
| Contig30890_pilon | 42474 | 357127 | Contig30890_pilon.2 | LG4 |
| Contig32501_pilon | 1 | 79400 | Contig32501_pilon.1 | LG8 |
| Contig32501_pilon | 81158 | 104315 | Contig32501_pilon.2 | LG5 |
| Contig34163_pilon | 1 | 276874 | Contig34163_pilon.1 | LG14 |
| Contig34163_pilon | 278116 | 477845 | Contig34163_pilon.2 | LG8 |
| Contig34793_pilon | 1 | 35174 | Contig34793_pilon.1 | LG13 |
| Contig34793_pilon | 35175 | 226445 | Contig34793_pilon.2 | LG4 |
| Contig37346_pilon | 1 | 181531 | Contig37346_pilon.1 | LG1 |
| Contig37346_pilon | 185444 | 510953 | Contig37346_pilon.2 | LG5 |
| Contig44873_pilon | 1 | 96939 | Contig44873_pilon.1 | LG3 |
| Contig44873_pilon | 100500 | 540225 | Contig44873_pilon.2 | LG2 |
| Contig53403_pilon | 1 | 162159 | Contig53403_pilon.1 | LG1 |
| Contig53403_pilon | 163594 | 231179 | Contig53403_pilon.2 | LG12 |
| Contig6264_pilon | 1 | 582129 | Contig6264_pilon.1 | LG1 |
| Contig6264_pilon | 582130 | 671930 | Contig6264_pilon.2 | LG16 |
| Contig6264_pilon | 675095 | 875693 | Contig6264_pilon.3 | LG1 |
| Contig67999_pilon | 1 | 75111 | Contig67999_pilon.1 | LG8 |
| Contig67999_pilon | 80918 | 230728 | Contig67999_pilon.2 | LG16 |
| Contig7355_pilon | 1 | 31398 | Contig7355_pilon.1 | LG4 |
| Contig7355_pilon | 31626 | 89218 | Contig7355_pilon.2 | LG7 |

**Extended Data Table 3.**
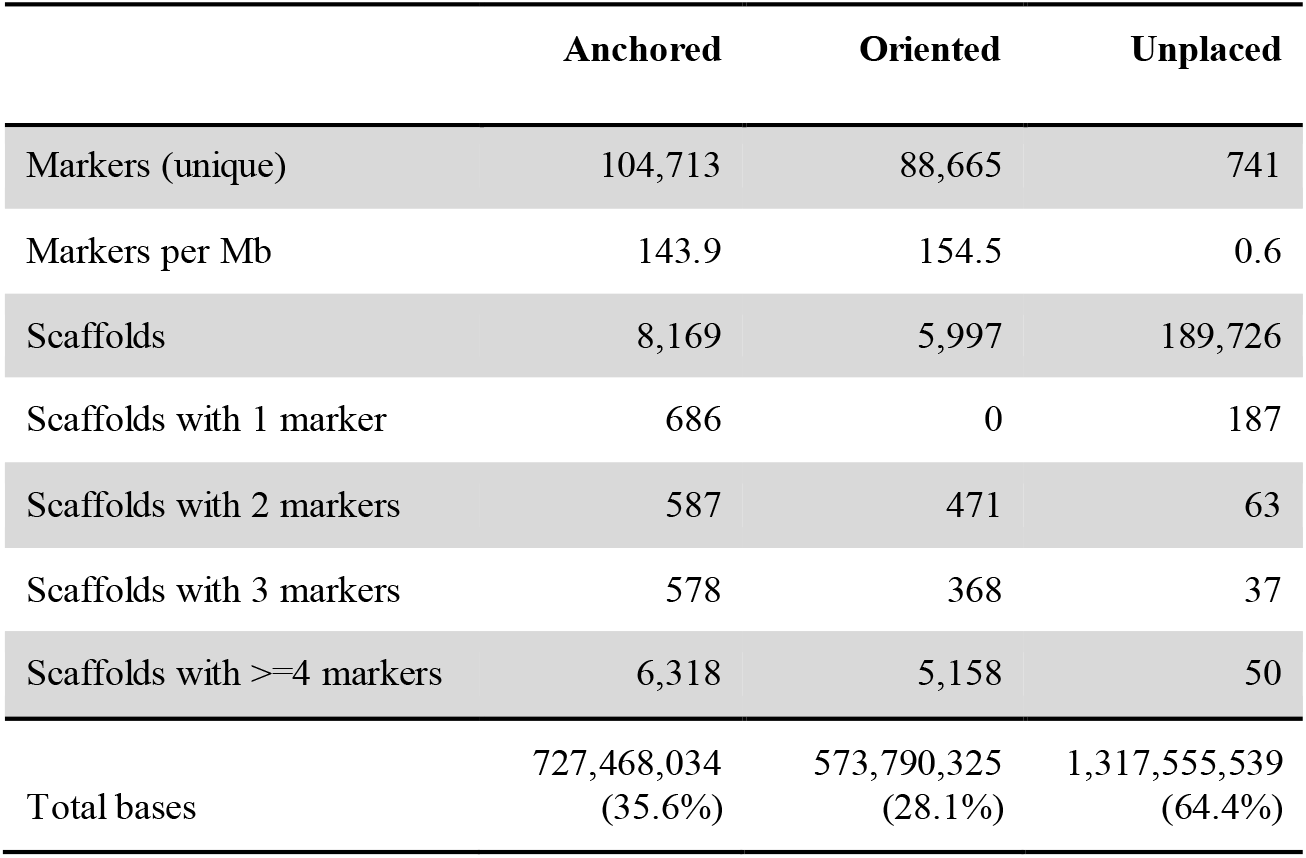
Summary statistics describing scaffold anchoring on *T. oceanicus* LOD5 linkage map markers

|  | Anchored | Oriented | Unplaced |
| --- | --- | --- | --- |
| Markers (unique) | 104,713 | 88,665 | 741 |
| Markers per Mb | 143.9 | 154.5 | 0.6 |
| Scaffolds | 8,169 | 5,997 | 189,726 |
| Scaffolds with 1 marker | 686 | 0 | 187 |
| Scaffolds with 2 markers | 587 | 471 | 63 |
| Scaffolds with 3 markers | 578 | 368 | 37 |
| Scaffolds with >=4 markers | 6,318 | 5,158 | 50 |
| Total bases | 727,468,034<br>(35.6%) | 573,790,325<br>(28.1%) | 1,317,555,539<br>(64.4%) |

**Extended Data Figure 3.**
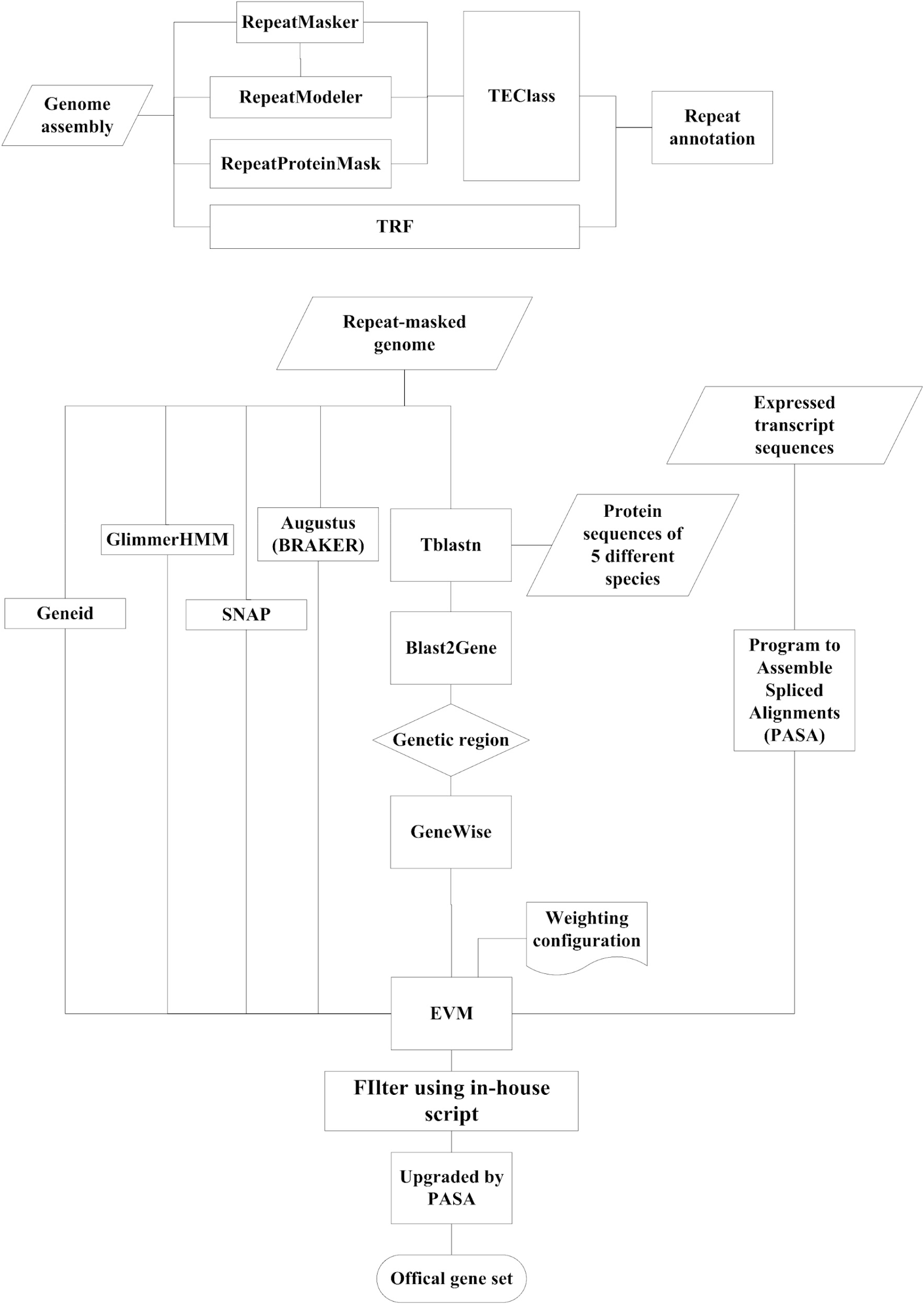
Workflow diagram of repeat annotation (top) and gene prediction (bottom) pipelines. Description of packages and parameters plus references are provided in the Methods section.

**Extended Data Table 4.**
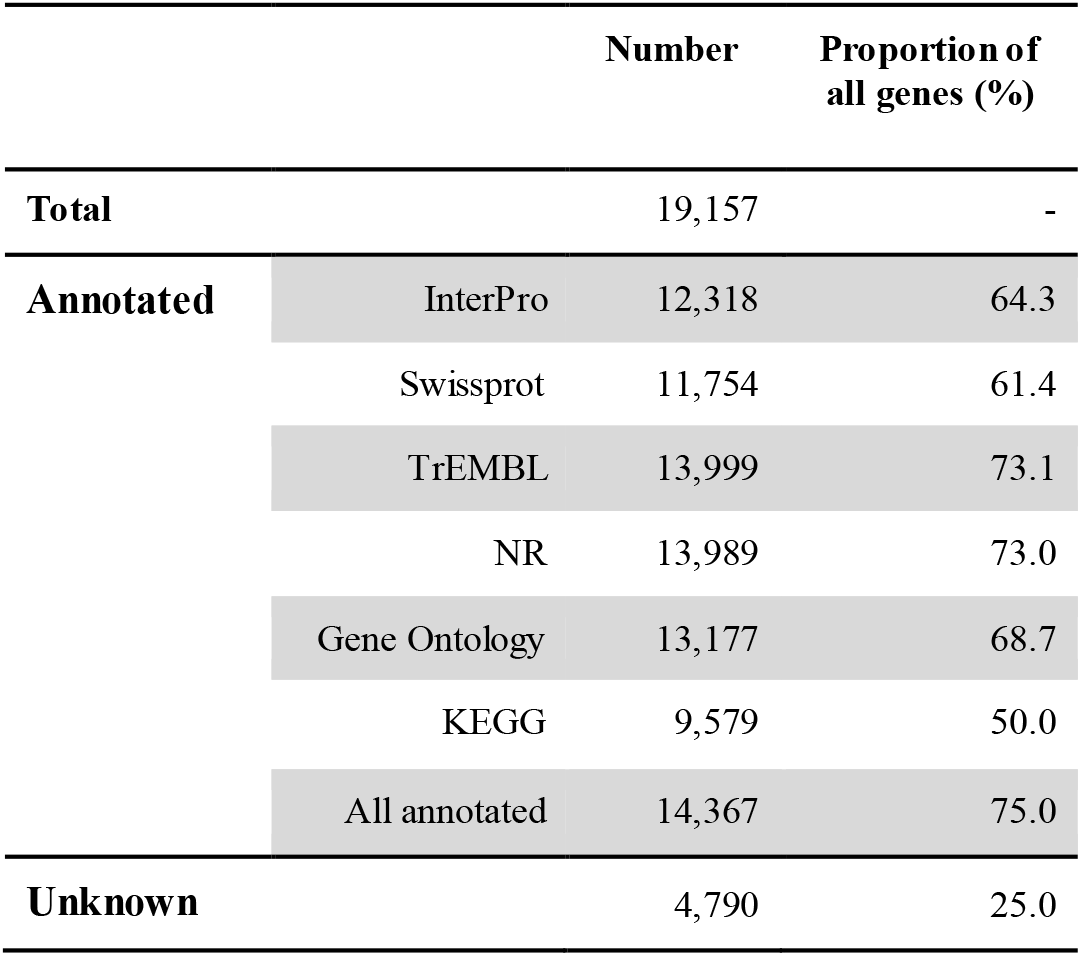
Functional annotation of *T. oceanicus* genes

|  |  | Number | Proportion of all genes (%) |
| --- | --- | --- | --- |
| **Total** |  | 19,157 | - |
| **Annotated** | InterPro | 12,318 | 64.3 |
|  | Swissprot | 11,754 | 61.4 |
|  | TrEMBL | 13,999 | 73.1 |
|  | NR | 13,989 | 73.0 |
|  | Gene Ontology | 13,177 | 68.7 |
|  | KEGG | 9,579 | 50.0 |
|  | All annotated | 14,367 | 75.0 |
| **Unknown** |  | 4,790 | 25.0 |

**Extended Data Figure 4.**
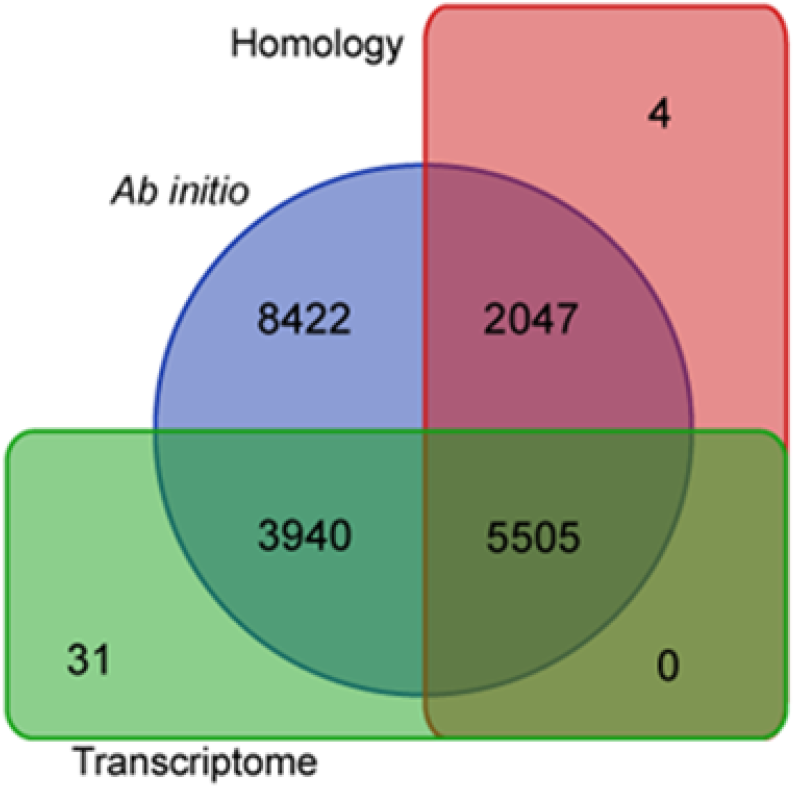
Venn diagram of genes predicted for *T. oceanicus* using different methods. Counts refer to the gene set that was obtained prior to final upgrade and filtering using PASA^71^, so the total gene number above is slightly higher than the final gene set. A detailed description of each pipeline is presented in the Methods.

**Extended Data Table 5.** Distribution of repetitive elements for each scaffolded *T. oceanicus* linkage group

| Linkage group | Transposable elements (kb) | % | Rank<sup>a</sup> | Tandem repeats (kb) | % | Rank<sup>a</sup> | Combined (kb) | % | Rank<sup>a</sup> |
| --- | --- | --- | --- | --- | --- | --- | --- | --- | --- |
| LG1 | 65992 | 38.6 | 8 | 1832 | 1.1 | 20 | 67823 | 39.7 | 11 |
| LG2 | 26518 | 35.7 | 17 | 1123 | 1.5 | 14 | 27641 | 37.2 | 17 |
| LG3 | 12908 | 36.3 | 14 | 572 | 1.6 | 13 | 13480 | 37.9 | 14 |
| LG4 | 16514 | 42.0 | 2 | 848 | 2.2 | 4 | 17362 | 44.2 | 2 |
| LG5 | 10253 | 36.2 | 15 | 500 | 1.8 | 11 | 10753 | 37.9 | 15 |
| LG6 | 23187 | 35.0 | 18 | 925 | 1.4 | 17 | 24113 | 36.4 | 18 |
| LG7 | 17533 | 36.3 | 13 | 936 | 1.9 | 8 | 18469 | 38.2 | 13 |
| LG8 | 12770 | 38.8 | 6 | 617 | 1.9 | 7 | 13387 | 40.7 | 6 |
| LG9 | 9952 | 38.2 | 9 | 659 | 2.5 | 2 | 10611 | 40.7 | 7 |
| LG10 | 9359 | 38.2 | 10 | 545 | 2.2 | 5 | 9904 | 40.5 | 8 |
| LG11 | 18920 | 34.1 | 19 | 804 | 1.5 | 15 | 19724 | 35.6 | 19 |
| LG12 | 4850 | 42.7 | 1 | 297 | 2.6 | 1 | 5148 | 45.4 | 1 |
| LG13 | 12684 | 37.4 | 12 | 624 | 1.8 | 10 | 13308 | 39.2 | 12 |
| LG14 | 12629 | 36.2 | 16 | 483 | 1.4 | 16 | 13112 | 37.6 | 16 |
| LG15 | 1298 | 41.7 | 3 | 53 | 1.7 | 12 | 1351 | 43.5 | 3 |
| LG16 | 6292 | 31.5 | 20 | 243 | 1.2 | 19 | 6535 | 32.7 | 20 |
| LG17 | 337 | 39.5 | 5 | 10 | 1.2 | 18 | 347 | 40.7 | 5 |
| LG18 | 699 | 38.2 | 11 | 42 | 2.3 | 3 | 741 | 40.5 | 9 |
| LG19 | 3 | 27.4 | 21 | 0 | 0.4 | 21 | 3 | 27.8 | 21 |
| Unplaced | 526597 | 40.0 | 4 | 25416 | 1.9 | 6 | 552013 | 41.9 | 4 |
| Total | 789295 | 38.6 | 7 | 36531 | 1.8 | 9 | 825826 | 40.4 | 10 |

<sup>a</sup> Sorted by the proportion of repetitive elements in linkage groups, from most to fewest

**Extended Data Figure 5.**
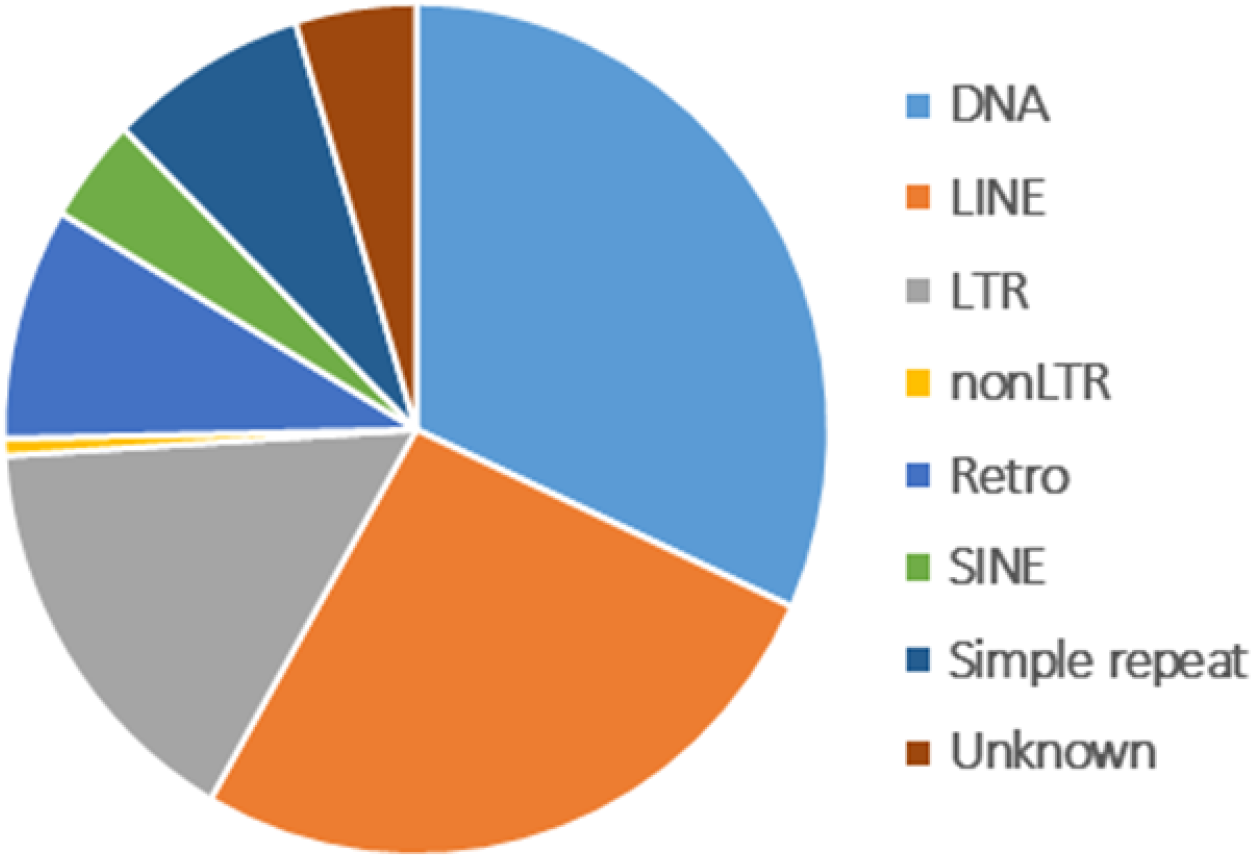
Proportions of eight major categories of transposable elements detected in the *T. oceanicus* genome. DNA = DNA transposons LTR = long terminal repeats LINE = long interspersed nuclear elements SINE = short interspersed nuclear elements Retro = retrotransposon

**Extended Data Table 6.** Comparison of gene features among ten insect species

| Species | Genome assembly size (Gb) | Gene number | Single exon gene number | % | Average gene length | Average CDS length (bp) | Average exon number per transcript | Average exon length (bp) | Ref. |
| --- | --- | --- | --- | --- | --- | --- | --- | --- | --- |
| **Field cricket** |  |  |  |  |  |  |  |  |  |
| *Teleogryllus oceanicus* | 2.0 | 19,157 | 1,266 | 6.6 | 12,232 | 1,184 | 6.2 | 395 | *this study* |
| **Migratory locust** |  |  |  |  |  |  |  |  |  |
| *Locusta migratoria* | 6.5 | 17,307 | 3,079 | 17.8 | 54,341<sup>a</sup> | 1,160 | 5.8 | 201 | <sup>62</sup> |
| **American Cockroach** |  |  |  |  |  |  |  |  |  |
| *Periplaneta americana* | 3.4 | 21,336 | 2,704 | 12.7 | 12,963<sup>b</sup> | - | 4<sup>b</sup> | 247<sup>b</sup> | <sup>94</sup> |
| **Fruit fly** |  |  |  |  |  |  |  |  |  |
| *Drosophila melanogaster* | 0.1 | 13,689 | 2,761 | 20.2 | 4,261 | 1,621 | 4.0 | 408 | <sup>64c</sup> |
| **Asian long-horned beetle** |  |  |  |  |  |  |  |  |  |
| *Anoplophora glabripennis* | 0.7 | 16,200 | 1,194 | 7.4 | 18,596 | 1,744 | 6.6 | 265 | <sup>65</sup> |
| **Bed bug** |  |  |  |  |  |  |  |  |  |
| *Cimex lectularius* | 0.7 | 13,751 | 1,410 | 10.3 | 29,076 | 1,846 | 10.2 | 265 | <sup>95</sup> |
| **Brown planthopper** |  |  |  |  |  |  |  |  |  |
| *Nilaparvata lugens* | 1.1 | 21,442 | 2,179 | 10.2 | 22,015 | 1,440 | 6.9 | 289 | <sup>66</sup> |
| **Dampwood termite** |  |  |  |  |  |  |  |  |  |
| *Zootermopsis nevadensis* | 0.5 | 15,129 | 370 | 2.4 | 24,927 | 1,890 | 9.4 | 365 | <sup>96</sup> |
| **Yellow fever mosquito** |  |  |  |  |  |  |  |  |  |
| *Aedes aegypti* | 1.3 | 19,585 | 1,158 | 5.9 | 36,583 | 2,144 | 6.4 | 481 | <sup>97</sup> |
| **Asian Tiger mosquito** |  |  |  |  |  |  |  |  |  |
| *Aedes albopictus* | 2.2 | 38,706 | 2,305 | 6.0 | 25,506 | 1,950 | 5.3 | 440 | <sup>98</sup> |

<sup>a</sup> Originally reported average transcript length
<sup>b</sup> Originally reported medians
<sup>c</sup> Adapted from<sup>63</sup>

**Extended Data Table 7.** Transposable element classification in *T. oceanicus* contrasted with three published genomes

|  | *T. oceanicus* |  | *L. migratoria*<sup>a</sup> |  | *D. melanogaster*<sup>a</sup> |  | *H. sapiens*<sup>a</sup> |  |
| --- | --- | --- | --- | --- | --- | --- | --- | --- |
| Types | Repeat size (bp) | % of genome | Repeat size (bp) | % of genome | Repeat size (bp) | % of genome | Repeat size (bp) | % of genome |
| DNA | 259181458 | 12.7 | 1,480,538,225 | 22.7 | 4,849,763 | 2.9 | 99,797,428 | 3.2 |
| LINE | 215705991 | 10.5 | 1,332,720,207 | 20.4 | 12,119,904 | 7.2 | 637,919,432 | 20.3 |
| LTR | 127951980 | 6.3 | 508,675,263 | 7.8 | 21,849,378 | 13.0 | 267,738,295 | 8.5 |
| nonLTR | 5233875 | 0.3 | 63,892,419 | 1.0 | - | - | - | - |
| Retro | 71828043 | 3.5 | 153,548,453 | 2.4 | - | - | - | - |
| SINE | 32344731 | 1.6 | 141,176,698 | 2.2 | 52,841 | 0.0 | 397,225,496 | 12.7 |
| Simple repeat | 63555524 | 3.1 | 13,026,240 | 0.2 | 2,733 | 0.0 | 26,240,511 | 0.8 |
| Unknown | 38615245 | 1.9 | 406,097,360 | 6.2 | 11,211,970 | 6.6 | 1,298,163 | 0.0 |
| Total | 789295269 | 38.6 | 3,840,808,141 | 58.9 | 50,785,143 | 30.0 | 1,434,373,137 | 46.0 |

DNA = DNA transposons
LTR = long terminal repeats
LINE = long interspersed nuclear elements
SINE = short interspersed nuclear elements
Retro = retrotransposon
<sup>a</sup> Data from<sup>61</sup>

**Extended Data Table 8.** Thoracic gene expression variation between embryonic crickets carrying *flatwing* vs. *normal-wing* genotypes

|  |  |
| --- | --- |
| Total number of genes passing expression filtering | 10,961 |
| Total DE genes | 830 |
| Up-regulated in *flatwing* | 328 |
| Down-regulated in *flatwing* | 502 |
| DE genes with log<sub>2</sub>FC > 1 | 204 |
| Up-regulated in *flatwing* | 25 |
| Down-regulated in *flatwing* | 179 |

**Extended Data Figure 6.**
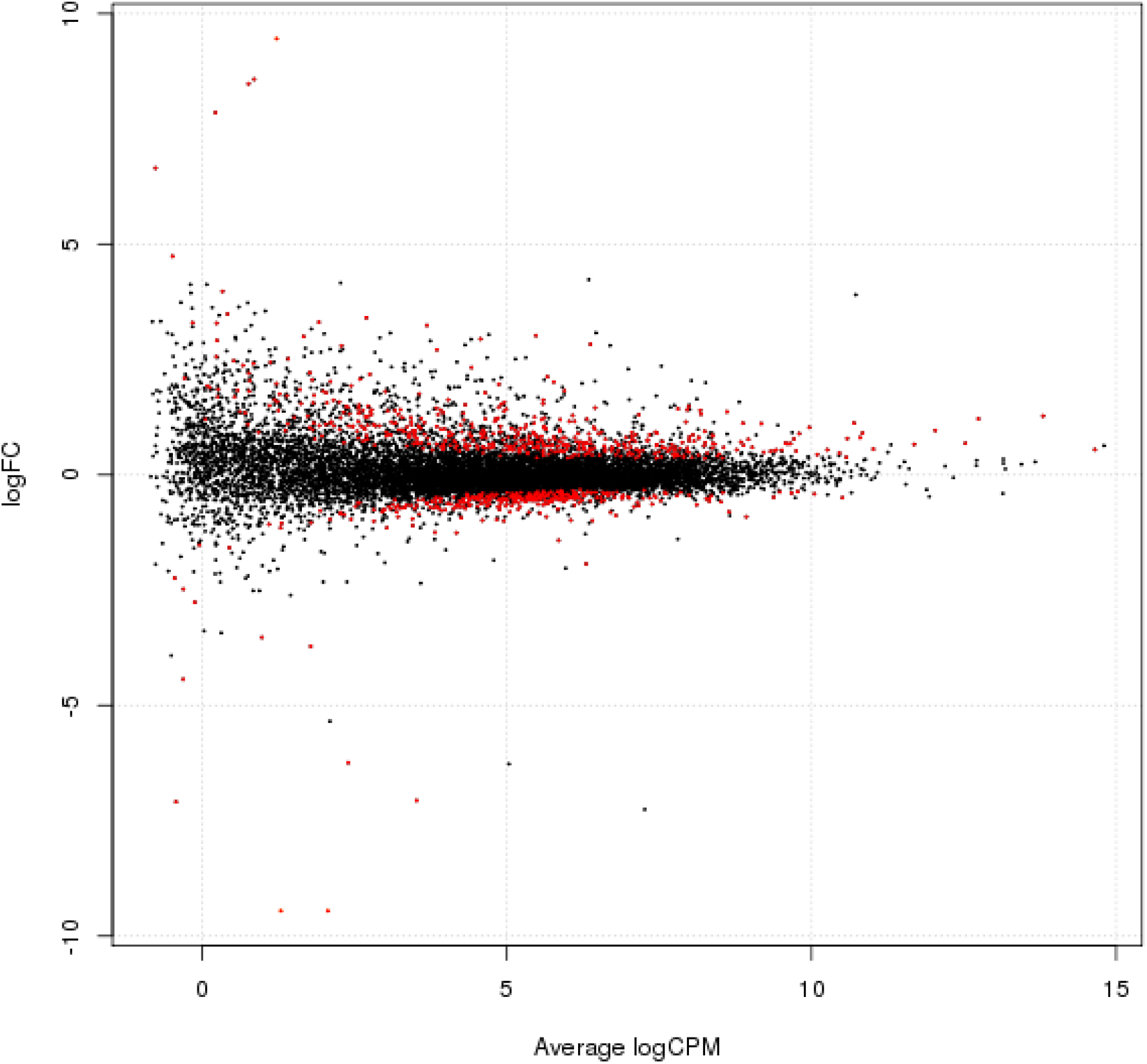
MA plot of thoracic genes DE between *T. oceanicus* embryos that were homozygous for *flatwing* vs. *normal-wing*. Red points indicate significantly differentially-expressed genes after filtering (see Methods), with positive values on the y-axis indicating genes downregulated in *flatwing* samples compared to *normal-wing* samples, and negative values indicating genes that are upregulated in *flatwing* samples. Log scales are base 2.

**Extended Data Table 9.** GO analysis of thoracic DEGs between embryos carrying *flatwing* vs. *normal-wing* genotypes

| GO | Type | Function | No. of DEGs | P-adj. |
| --- | --- | --- | --- | --- |
| GO:0003824 | molecular_function | catalytic activity | 216 | 0.006 |
| GO:0016787 | molecular_function | hydrolase activity | 80 | 0.027 |
| GO:0044087 | biological_process | regulation of cellular component biogenesis | 20 | 0.004 |
| GO:0051493 | biological_process | regulation of cytoskeleton organization | 12 | 0.023 |
| GO:0090066 | biological_process | regulation of anatomical structure size | 12 | 0.009 |
| GO:0097435 | biological_process | supramolecular fiber organization | 12 | 0.023 |
| GO:0032535 | biological_process | regulation of cellular component size | 11 | 0.004 |
| GO:0032956 | biological_process | regulation of actin cytoskeleton organization | 11 | 0.001 |
| GO:0032970 | biological_process | regulation of actin filament-based process | 11 | 0.001 |
| GO:1902903 | biological_process | regulation of supramolecular fiber organization | 11 | 0.002 |
| GO:0043254 | biological_process | regulation of protein complex assembly | 10 | 0.035 |
| GO:0110053 | biological_process | regulation of actin filament organization | 10 | 0.001 |
| GO:0008064 | biological_process | regulation of actin polymerization or depolymerization | 8 | 0.006 |
| GO:0030832 | biological_process | regulation of actin filament length | 8 | 0.006 |
| GO:0030833 | biological_process | regulation of actin filament polymerization | 8 | 0.006 |
| GO:0032271 | biological_process | regulation of protein polymerization | 8 | 0.015 |
| GO:0015630 | cellular_component | microtubule cytoskeleton | 6 | 0.001 |
| GO:0005248 | molecular_function | voltage-gated sodium channel activity | 5 | 0.004 |
| GO:0010927 | biological_process | cellular component assembly involved in morphogenesis | 5 | 0.010 |
| GO:0034706 | cellular_component | sodium channel complex | 4 | 0.007 |

**Extended Data Table 10.**
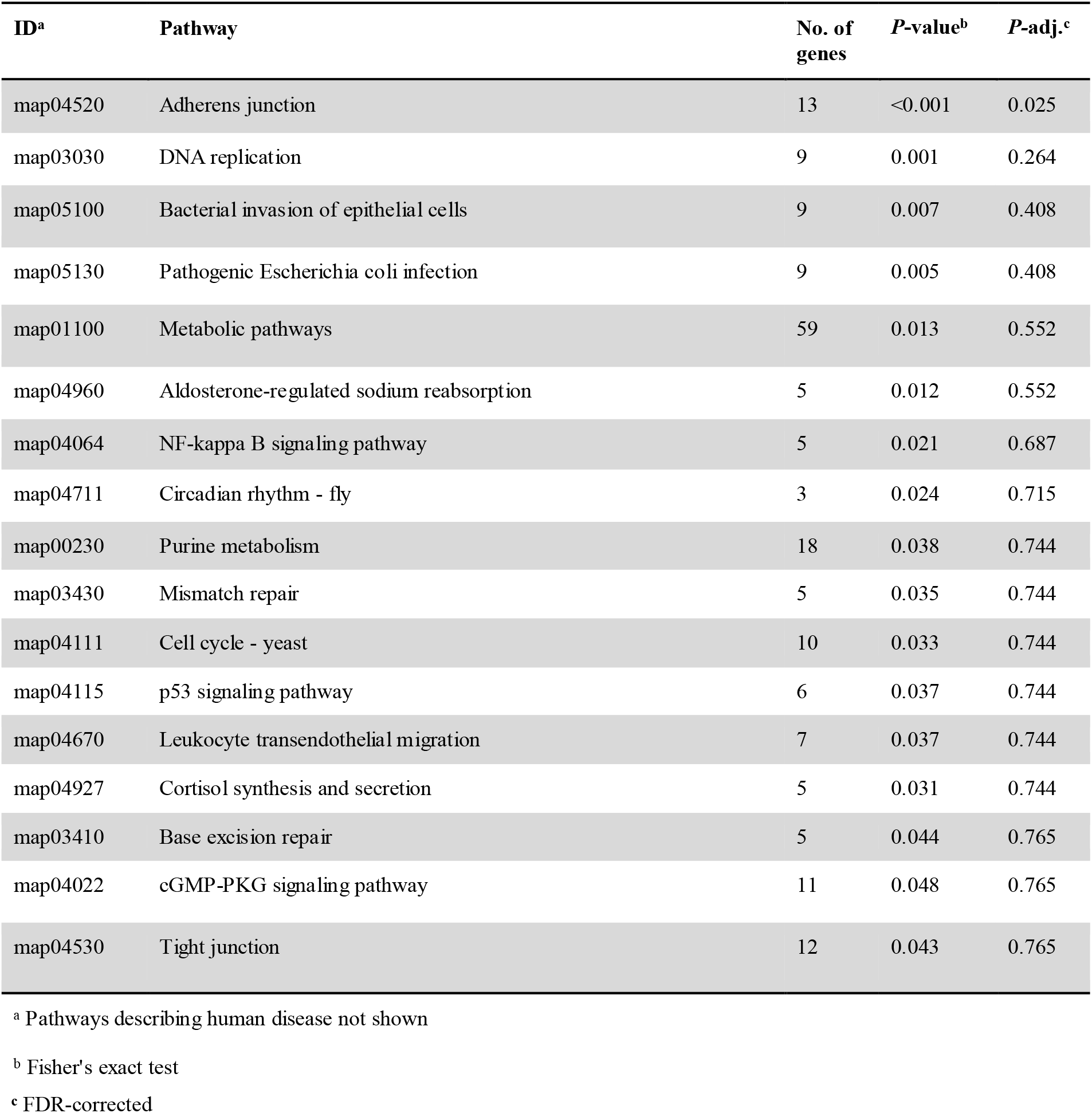
KEGG pathway enrichment of thoracic DEGs between embryos carrying *flatwing* vs. *normal-wing* genotypes

**Extended Data Table 11.** Top candidate genes associated with flatwing

| Coordinates |  |  |  |  |  |  |
| --- | --- | --- | --- | --- | --- | --- |
| Scaffold | Start | Stop | Gene ID |  | Description<sup>a</sup> | Source<sup>b</sup> |
| Contig10220_pilon | 78887 | 163134 | TOT000182.1 | YGD6 | Zinc-type alcohol dehydrogenase-like protein C1773.06c | Top 1% |
| Contig11287_pilon | 199864 | 215783 | TOT001075.1 | HNF4 | Transcription factor HNF-4 homolog | Top 1% |
| Contig12423_pilon | 12805 | 162659 | TOT001854.1 | OSM3 | Osmotic avoidance abnormal protein 3 | Top 1% |
| Contig12752_pilon | 14326 | 68612 | TOT002129.1 | ROR1 | Inactive tyrosine-protein kinase transmembrane receptor ROR1 | Top 1% |
| Contig12919_pilon | 275064 | 329308 | TOT002239.1 | CLOCK | Circadian locomotor output cycles protein kaput | Top 1%<br>DEG |
| Contig13810_pilon | 43734 | 77412 | TOT002877.1 | CRTAP | Cartilage-associated protein | Top 1% |
| Contig140_pilon | 14528 | 100227 | TOT003072.1 | PP4R1 | Serine/threonine-protein phosphatase 4 regulatory subunit 1 | Top 1% |
| Contig17198_pilon | 217974 | 328434 | TOT004721.1 | SCN60 | Sodium channel protein 60E | Top 1% |
| Contig17198_pilon | 398116 | 526355 | TOT004722.1 | SCN60 | Sodium channel protein 60E | Top 1% |
| Contig17528_pilon | 151965 | 161397 | TOT004867.1 | OBSTE | Protein obstructor-E | Kauai<br>DEG |
| Contig17791_pilon | 294998 | 303418 | TOT005017.1 | PKRA | Protein krasavietz | DEG<br>QTL |
| Contig18506_pilon | 5030 | 106086 | TOT005335.1 | STRN3 | Striatin-3 | Top 1% |
| Contig20777_pilon | 197732 | 433721 | TOT006213.1 | COLL | Transcription factor collier | Top 1%<br>BSA |
| Contig23358_pilon | 266813 | 357642 | TOT006927.1 | E78C | Ecdysone-induced protein 78C (Eip78C) | Top 1% |
| Contig23647_pilon | 61437 | 289082 | TOT006991.1 | RAVR1 | Ribonucleoprotein PTB-binding 1 | Top 1% |
| Contig24519_pilon | 221508 | 332371 | TOT007217.1 | A0A167<br>WTZ1 | Endo-1,3(4)-beta-glucanase | Top 1% |
| Contig24519_pilon | 569981 | 635619 | TOT007221.1 | SEPIA | Pyrimidodiazepine synthase | Top 1% |
| Contig30320_pilon | 33122 | 79411 | TOT008755.1 | PTPC1 | Protein tyrosine phosphatase domain-containing protein 1 | Top 1% |
| Contig3077_pilon | 487713 | 492969 | TOT008894.1 | REXO4 | RNA exonuclease 4 | Top 1% |
| Contig31374_pilon | 378769 | 413960 | TOT009065.1 | CPT2 | Carnitine O-palmitoyltransferase 2, mitochondrial | Top 1% |
| Contig31374_pilon | 461061 | 489320 | TOT009067.1 | FRM4B | FERM domain-containing protein 4B | Top 1% |
| Contig32190_pilon | 94344 | 248306 | TOT009274.1 | RN207 | RING finger protein 207 | Top 1% |
| Contig35402_pilon | 14084 | 125884 | TOT010060.1 | ABCG1 | ATP-binding cassette sub-family G member 1 | Top 1% |
| Contig37346_pilon.1 | 133394 | 180067 | TOT010542.1 | SCYL1 | N-terminal kinase-like protein | Top 1% |
| Contig40107_pilon | 150347 | 172207 | TOT011176.1 | THUM3 | THUMP domain-containing protein 3 | DEG,<br>QTL |
| Contig4430_pilon | 60074 | 108676 | TOT012009.1 | LAR | Tyrosine-protein phosphatase Lar | Top 1% |
| Contig4497_pilon | 323 | 114981 | TOT012126.1 | MYO5A | Unconventional myosin-Va | Top 1% |
| Contig48084_pilon | 4534 | 15580 | TOT012711.1 | KPEL | Serine/threonine-protein kinase pelle | Top 1% |
| Contig48322_pilon | 73569 | 78934 | TOT012764.1 | CAH10 | Carbonic anhydrase-related protein 10 | Top 1% |
| Contig52923_pilon | 4299 | 134158 | TOT013504.1 | RENT2 | Regulator of nonsense transcripts 2 | Top 1%<br>DEG |
| Contig52923_pilon | 172817 | 234071 | TOT013505.1 | NBL1 | Neuroblastoma suppressor of tumorigenicity 1 | Top 1% |
| Contig53931_pilon | 135337 | 200203 | TOT013689.1 | TADBP | TAR DNA-binding protein 43 | Top 1% |
| Contig55532_pilon | 2641 | 6823 | TOT013967.1 | SOSSC | SOSSC_BOVIN | Top 1% |
| Contig5817_pilon | 13001 | 94458 | TOTO14395.1 | A0A1B6<br>LWD6 | Uncharacterized protein | Top 1% |
| Contig6025_pilon | 181847 | 338853 | TOT014693.1 | PAX6 | Paired box protein Pax-6 | Top 1% |
| Contig6181_pilon | 7490 | 15461 | TOT014894.1 | MYCT | Proton myo-inositol cotransporter | Top 1% |
| Contig6371_pilon | 72321 | 126646 | TOT015146.5 | GOGA4 | Golgin subfamily A member 4 | Top 1% |
| Contig6636_pilon | 248427 | 279756 | TOT015511.1 | A0A067<br>RPQ2 | LRR domain-containing protein | Top 1% |
| Contig6636_pilon | 332473 | 344815 | TOT015512.1 | IPR0110<br>11 | Uncharacterized protein | Top 1% |
| Contig66512_pilon | 19778 | 188429 | TOT015537.1 | LASP1 | LIM and SH3 domain protein F42H10.3 | Top 1%<br>DEG |
| Contig6932_pilon | 94582 | 114944 | TOT015868.1 | ABCB8 | ATP-binding cassette sub-family B member 8,<br>mitochondrial | Top 1% |
| Contig6932_pilon | 132941 | 150338 | TOT015869.1 | APMAP | Adipocyte plasma membrane-associated protein | Top 1% |
| Contig7020_pilon | 57223 | 96643 | TOT015999.1 | SPS1 | Selenide, water dikinase | Top 1% |
| Contig7210_pilon | 172512 | 326460 | TOT016305.1 | MYO | myoglianin | Top 1% |
| Contig7490_pilon | 12720 | 16839 | TOT016652.1 | GCN5 | Histone acetyltransferase GCN5 | Top 1% |
| Contig8190_pilon | 192630 | 256540 | TOT017431.1 | AT133 | Probable cation-transporting ATPase 13A3 | Top 1% |
| Contig82459_pilon | 106133 | 176987 | TOT017512.1 | UNC89 | Muscle M-line assembly protein unc-89 | DEG<br>QTL |
| Contig83863_pilon | 1777 | 51856 | TOT017662.1 | A0A017<br>RSC4 | Uncharacterized protein | Top 1% |
| Contig92683_pilon | 43257 | 76189 | TOT018508.1 | A0A0T6<br>B8G7 | Uncharacterized protein | Top 1% |
| Contig33215_pilon | 67326 | 419738 | TOT009518.1 | PLXA4 | Plexin-A4 | QTL<br>BSA |
| Contig43580_pilon | 106377 | 137341 | TOT011864.1 | RNF41 | E3 ubiquitin-protein ligase NRDP1 | QTL<br>BSA |

<sup>a</sup> Functional descriptions and references provided in Main Text
<sup>b</sup> The criterion for inclusion as a top candidate was that a gene had to receive support for association with the flatwing phenotype from at least two of the following four sources:
QTL = portion of the gene was located within a 1 kb flanking region of a significantly-associated (FDR-corrected) marker in the flatwing QTL analysis
Top1% = portion of the gene was located within a 1 kb flanking region of a SNP in the top 1% of significantly-associated markers in the flatwing QTL analysis (Top1% candidates automatically received “QTL” support)
BSA = portion of the gene was located within a 1 kb flanking region of a significantly-associated marker from the previously-published bulked segregant analysis of Kauai flatwing males<sup>7</sup> which was also anchored to LG1
DEG = gene was significantly differentially expressed between pure-breeding *normal-wing* genotypes and *flatwing* genotypes, in embryonic thoracic tissue (track iv of Fig. 2a in Main Text)

**Extended Data Table 12.** GO analysis of candidate flatwing-associated genes (CGs)

| GO | Type | Function | No. of CGs | *P*-adj. ($\chi^2$ test) |
| --- | --- | --- | --- | --- |
| GO:0010720 | biological_process | positive regulation of cell development | 3 | <0.001 |
| GO:0045597 | biological_process | positive regulation of cell differentiation | 3 | 0.002 |
| GO:0060284 | biological_process | regulation of cell development | 3 | 0.042 |
| GO:0003707 | molecular_function | steroid hormone receptor activity | 2 | <0.001 |
| GO:0009755 | biological_process | hormone-mediated signaling pathway | 2 | 0.006 |
| GO:0030545 | molecular_function | receptor regulator activity | 2 | 0.035 |
| GO:0043401 | biological_process | steroid hormone mediated signaling pathway | 2 | 0.002 |
| GO:0045666 | biological_process | positive regulation of neuron differentiation | 2 | 0.040 |
| GO:0048018 | molecular_function | receptor ligand activity | 2 | 0.022 |

**Extended Data Table 13.** Identification of *Teleogryllus oceanicus* cuticular hydrocarbon profile peaks using gas chromatography - mass spectrometry

| Peak<sup>a</sup> | Kováts retention index<sup>b</sup> | Identification | Diagnostic ions |
| --- | --- | --- | --- |
| standard |  | pentadecane |  |
| 1 | 2840 | 4MeC<sub>28</sub> | 365, 71 |
| 2 | 2893 | 10MeC<sub>28</sub> | 281, 155 |
| 3 | 2910 | 13MeC<sub>29</sub> | 252, 196 |
| 4 | 3028 | C<sub>30:1</sub> | 434 |
| 5 | 3043 | 4MeC<sub>30</sub> | 436, 393, 71 |
| 6 | 3064 | 7,x-diMeC<sub>30</sub> | 365, 112 |
| 7 | 3075 | unidentified |  |
| 8 | 3086 | C<sub>31:1</sub> | 434 |
| 9 | 3110 | 11MeC<sub>31</sub> | 308, 168 |
| 10 | 3119 | 7,8MeC<sub>31</sub> | 364, 112 |
| 11 | 3130 | 7-C<sub>31</sub>ene | 434, 528<sup>c</sup>, 145<sup>c</sup>, 383<sup>c</sup> |
| 12 | 3142 | C<sub>31:1</sub> | 434 |
| 13 | 3152 | C<sub>31:2</sub> | 432 |
| 14 | 3161 | C<sub>31:2</sub> | 432 |
| 15 | 3174 | C<sub>31:2</sub> | 432 |
| 16 | 3188 | C<sub>31:2</sub> | 432 |
| 17 | 3242 | 4MeC<sub>32</sub> | 421, 71 |
| 18 | 3252 | unidentified |  |
| 19 | 3268 | C<sub>32:2</sub> | 446 |
| 20 | 3288 | C<sub>33:1</sub> | 462 |
| 21 | 3331 | C<sub>33:1</sub> | 462 |
| 22 | 3347 | C<sub>33:2</sub> | 460 |
| 23 | 3355 | C<sub>33:2</sub> | 460 |
| 24 | 3365 | C<sub>33:2</sub> | 460 |
| 25 | 3379 | 3,x-C<sub>33:2</sub> | 460, 647<sup>c</sup>, 89<sup>c</sup> |
| 26 | 3391 | C<sub>33:2</sub> | 460 |

<sup>a</sup> Peak identification is based on Table S4 of Pascoal et al. (2016)<sup>83</sup>, reproduced here. The 26 quantified peaks are presented in sequential order in the table and in Fig. 3A of the main text.
<sup>b</sup> Calculation of the Kováts retention index using *n*-alkane standards (C<sub>7</sub> - C<sub>40</sub>) is described in Majlát et al. (1974)<sup>99</sup>
<sup>c</sup> Ions used when identifying with dimethyl-disulphide derivation

**Extended Data Table 14.** Principal components with eigenvalues > 1 from PCA on male CHC profiles; MANOVA results examine the effect of male morph on scores for each PC (multivariate model: Wilks’ λ = 0.517, F_6,191_ = 29.769, p < 0.001).

| Principal component | Eigenvalue | % Variance explained | $F_{1,196}$ | P | $R^2$ |
| --- | --- | --- | --- | --- | --- |
| 1 | 9.408 | 36.18 | 25.885 | <0.001 | 0.131 |
| 2 | 2.635 | 10.136 | 18.040 | <0.001 | 0.092 |
| 3 | 2.490 | 9.576 | 21.454 | <0.001 | 0.109 |
| 4 | 1.888 | 7.261 | 0.001 | 0.979 | 0.000 |
| 5 | 1.315 | 5.058 | 25.741 | <0.001 | 0.131 |
| 6 | 1.020 | 3.925 | 4.079 | 0.043 | 0.021 |

**Extended Data Figure 7.**
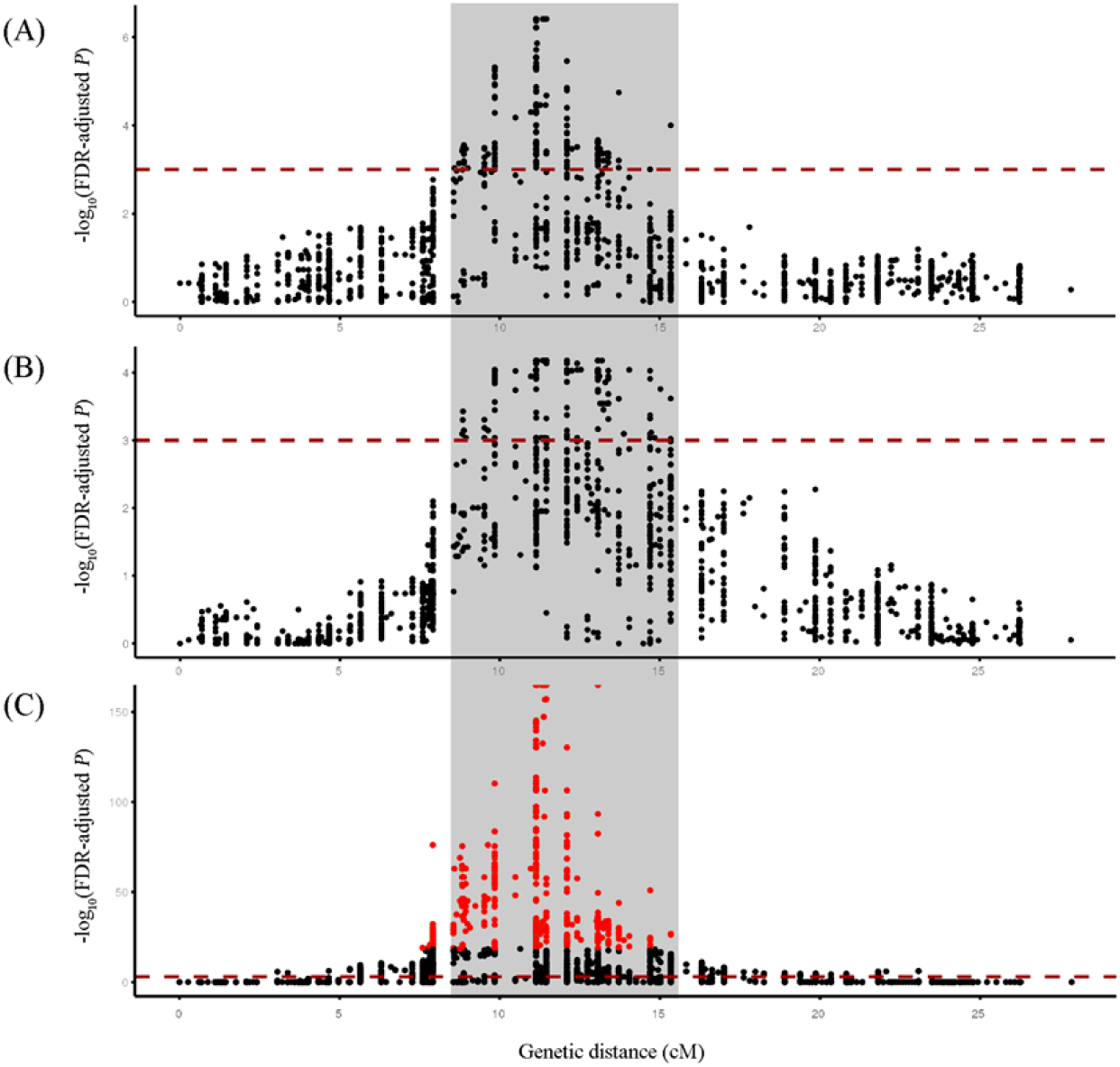
Genomic regions associated with different principal components describing male CHC profiles. Manhattan plot for LG1 (putative X chromosome) showing **a,** the PC4-associated QTL, **b,** PC6-associated QTL and **c,** the *flatwing* QTL for comparison. The horizontal dashed lines indicate FDR-corrected significance threshold of P < 0.001, and the top 1% most significant flatwing-associated QTL markers are plotted in red in **c**. The light grey rectangle spans the genetic region in which flatwing-associated markers and CHC principal component-associated markers colocalize.

**Extended Data Table 15.**
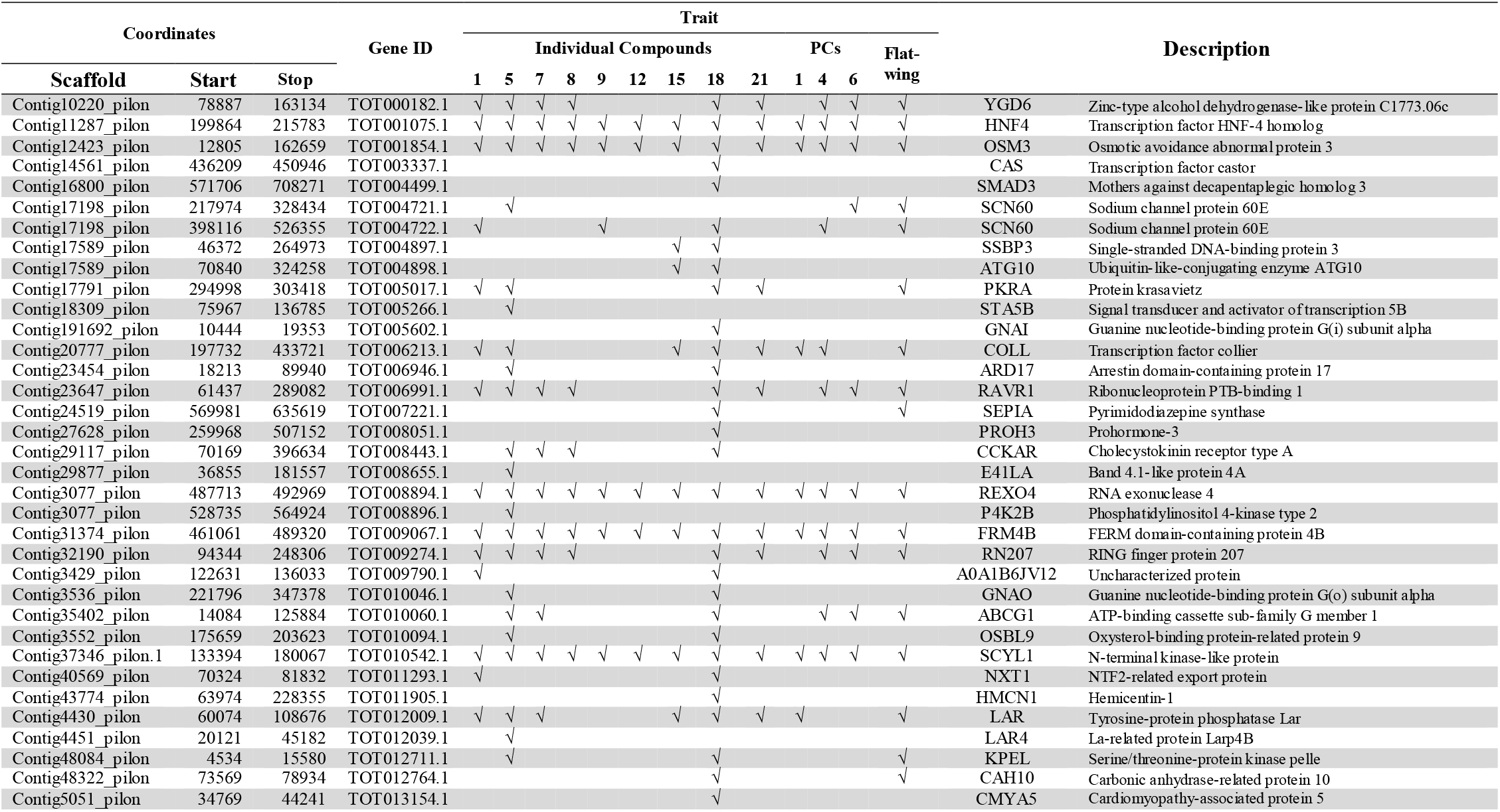
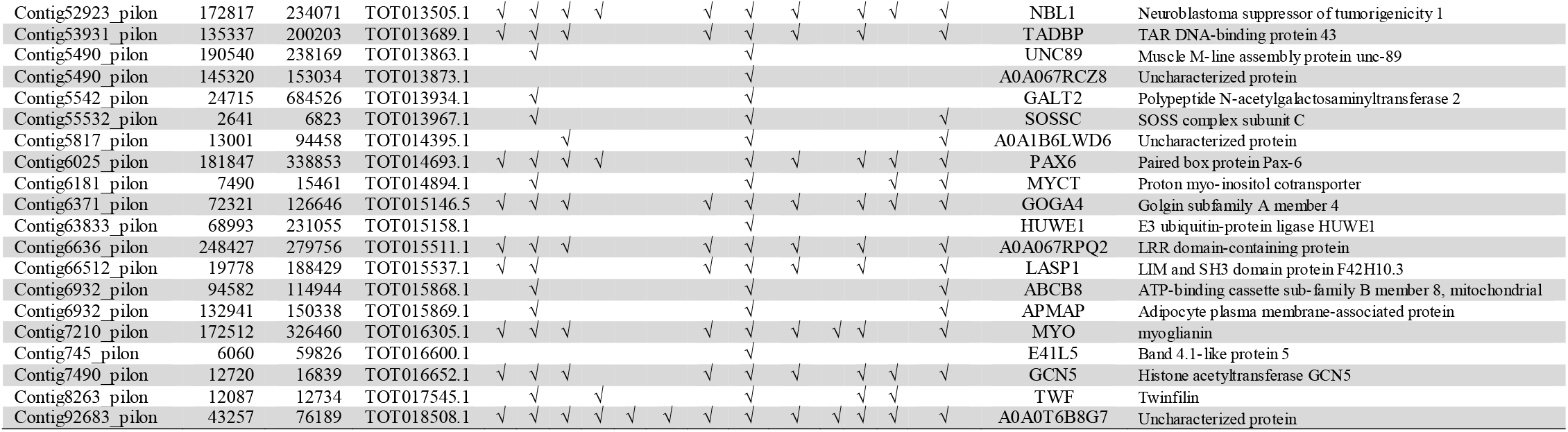
Candidate gene set associated with each CHC phenotype (individual or principal component) that yielded a significant QTL on the putative X (LG1), with the flatwing QTL for comparison

**Extended Data Table 16.** Allele replacement table for identifying the X chromosome in the *T. oceanicus* linkage map

| **Dam** | **Sire** | **Sire**<br>**assumed** | **Sire**<br>**rewrite** | **male**<br>**current** | **male**<br>**rewrite** | **male**<br>**current** | **male**<br>**rewrite** |
| --- | --- | --- | --- | --- | --- | --- | --- |
| **0/1** | 0/0 | 0/. | 0/2 | 0/0 | 0/2 | 1/1 | 1/2 |
| **0/1** | 1/1 | 1/. | 1/2 | 0/0 | 0/2 | 1/1 | 1/2 |
| **0/1** | 2/2 | 2/. | NA | - | - | - | - |
| **0/2** | 0/0 | 0/. | 0/1 | 0/0 | 0/1 | 2/2 | 1/2 |
| **0/2** | 1/1 | 1/. | NA | - | - | - | - |
| **0/2** | 2/2 | 2/. | 2/1 | 0/0 | 0/1 | 2/2 | 2/1 |
| **1/2** | 0/0 | 0/. | NA | - | - | - | - |
| **1/2** | 1/1 | 1/. | 1/3 | 1/1 | 1/3 | 2/2 | 2/3 |
| **1/2** | 2/2 | 2/. | 2/3 | 1/1 | 1/3 | 2/2 | 2/3 |

**Extended Data Figure 8.**
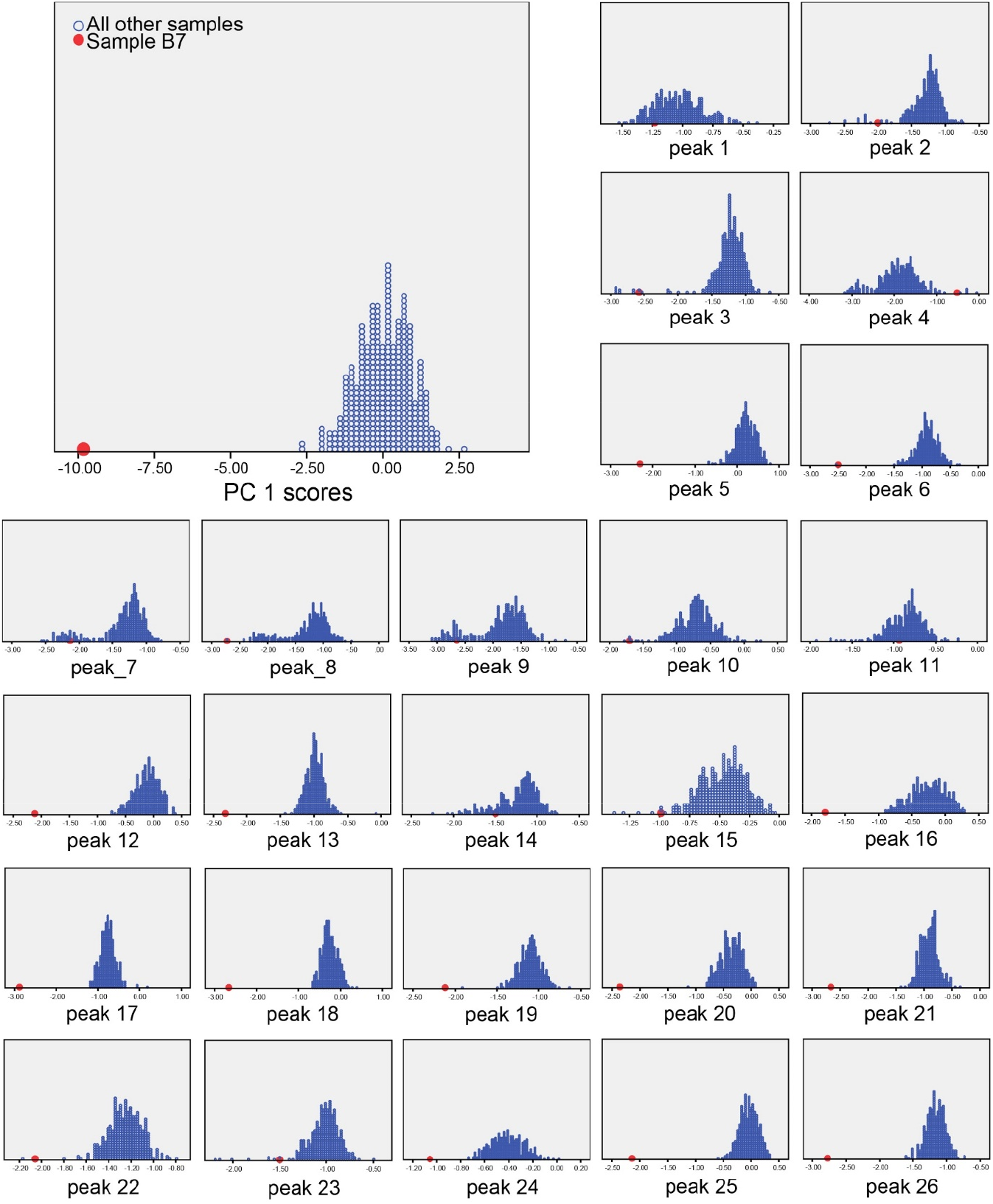
Histograms illustrating the identification of a CHC sample outlier. Sample B7, a normal-wing male, is indicated by the enlarged red dot in each plot. The sample was observed on visual inspection to deviate substantially from the distribution of principal component 1 scores for all other mapping individuals. Further inspection revealed this also to be the case in the majority of cases when the sample was assessed for each CHC peak individually. It was thus excluded from further analysis.

## Acknowledgements

The Natural Environment Research Council provided funding to N.W.B. (NE/G014906/1, NE/L011255/1), and to N.W.B. and M.G.R. (NE/1027800/1). Sequencing support was provided by Edinburgh Genomics and the Centre for Genomic Research at the University of Liverpool. Bioinformatics resources at St Andrews were funded by a Wellcome Trust ISSF award (105621/Z/14/Z). Support from the China Scholarship Council (201703780018) to X.Z. is gratefully acknowledged. The Biotechnology and Biological Sciences Research Council provided support to M.B. which aided in the development of ChirpBase (BB/K020161/1). We thank J. Kenny for NGS sequencing advice; Y. Fang for read processing; R. Fallon for bioinformatics assistance; J.Q. Liu and K. Wang for advice regarding gene prediction pipelines; J. Bastiaansen and P. Gienapp for assistance with ASReml, W.V. Bailey, B. Gray, J.T. Rotenberry, S. Vardy and M. Zuk for assistance in the field; D. Forbes, A. Grant and T. Sneddon for assistance in the laboratory; C. Mitchell for assistance with CHC analyses.

## Author contributions

N.W.B. conceived and led the study. S.P., K.G., M.B., M.G.R. and N.W.B. designed experiments. S.P. led data collection. S.P. did genetic crosses and wet lab work. S.P., J.E.R., X.Z., T.C., E.L., X.L., J.H., J.G.R., B.L.S., U.T. and N.W.B. performed analyses. M.B., R.J.C., S.J., E.L., M.B. and N.W.B. designed ChirpBase. N.W.B. led manuscript writing. S.P., J.E.R., X.Z., E.L., M.B., M.G.R. and N.W.B contributed to writing.

## References

1. Wiens, J. J. Widespread loss of sexually selected traits: how the peacock lost its spots. Trends in Ecology & Evolution 16, 517–523 (2001).

2. Whiting, M. F., Bradler, S. & Maxwell, T. Loss and recovery of wings in stick insects. Nature 421, 264–267 (2003).

3. Nadeau, J. H. et al. Pleiotropy, homeostasis, and functional networks based on assays of cardiovascular traits in genetically randomized populations. Genome Res 13, 2082–2091 (2003).

4. Zuk, M., Rotenberry, J. T. & Tinghitella, R. M. Silent night: adaptive disappearance of a sexual signal in a parasitized population of field crickets. Biol Lett 2, 521–524 (2006).

5. Zuk, M., Bailey, N. W., Gray, B. & Rotenberry, J. T. Sexual signal loss: The link between behaviour and rapid evolutionary dynamics in a field cricket. J Anim Ecol 87, 623–633 (2018).

6. Tinghitella, R. M. Rapid evolutionary change in a sexual signal: genetic control of the mutation ‘flatwing’ that renders male field crickets (Teleogryllus oceanicus) mute. Heredity 100, 261–267 (2008).

7. Pascoal, S. et al. Rapid convergent evolution in wild crickets. Curr Biol 24, 1369–1374 (2014).

8. Fisher, R. The Genetical Theory of Natural Selection. 1st edn, (The Clarendon Press, 1930).

9. Orr, H. A. The genetic theory of adaptation: a brief history. Nat Rev Genet 6, 119–127 (2005).

10. Hartley, C. J. et al. Amplification of DNA from preserved specimens shows blowflies were preadapted for the rapid evolution of insecticide resistance. Proc Natl Acad Sci U S A 103, 8757–8762 (2006).

11. Raymond, M., Berticat, C., Weill, M., Pasteur, N. & Chevillon, C. Insecticide resistance in the mosquito *Culex pipiens*: what have we learned about adaptation? Genetica 112-113, 287–296 (2001).

12. van’t Hof, A. E., Edmonds, N., Dalikova, M., Marec, F. & Saccheri, I. J. Industrial melanism in British peppered moths has a singular and recent mutational origin. Science 332, 958–960 (2011).

13. Van’t Hof, A. E. et al. The industrial melanism mutation in British peppered moths is a transposable element. Nature 534, 102–105 (2016).

14. True, J. R. Insect melanism: the molecules matter. Trends in Ecology & Evolution 18, 640–647 (2003).

15. Challis, R. J., Kumar, S., Stevens, L. & Blaxter, M. GenomeHubs: simple containerized setup of a custom Ensembl database and web server for any species. Database (Oxford) 2017, doi:10.1093/database/bax039 (2017).

16. Snodgrass, R. E. Principles of Insect Morphology (Cornell University Press, 1993).

17. De Celis, J. F. Pattern formation in the *Drosophila* wing: The development of the veins. Bioessays 25, 443–451 (2003).

18. Hatini, V., Kula-Eversole, E., Nusinow, D. & Del Signore, S. J. Essential roles for stat92E in expanding and patterning the proximodistal axis of the *Drosophila* wing imaginal disc. Dev Biol 378, 38–50 (2013).

19. Vervoort, M., Crozatier, M., Valle, D. & Vincent, A. The COE transcription factor Collier is a mediator of short-range Hedgehog-induced patterning of the *Drosophila* wing. Curr Biol 9, 632–639 (1999).

20. Hevia, C. F. & de Celis, J. F. Activation and function of TGFbeta signalling during *Drosophila* wing development and its interactions with the BMP pathway. Dev Biol 377, 138–153 (2013).

21. Bicocca, V. T. et al. Crosstalk between ROR1 and the Pre-B cell receptor promotes survival of t(1;19) acute lymphoblastic leukemia. Cancer Cell 22, 656–667 (2012).

22. Carvajal-Gonzalez, J. M., Mulero-Navarro, S., Smith, M. & Mlodzik, M. A novel Frizzled-based screening tool identifies genetic modifiers of planar cell polarity in *Drosophila* wings. G3 (Bethesda) 6, 3963–3973 (2016).

23. Adler, P. N. The frizzled/stan pathway and planar cell polarity in the *Drosophila* wing. Curr Top Dev Biol 101, 1–31 (2012).

24. Wu, C. et al. Pelle modulates dFoxO-mediated cell death in *Drosophila*. Plos Genet 11, e1005589, doi:10.1371/journal.pgen.1005589 (2015).

25. Carre, C., Szymczak, D., Pidoux, J. & Antoniewski, C. The histone H3 acetylase dGcn5 is a key player in *Drosophila melanogaster* metamorphosis. Mol Cell Biol 25, 8228–8238 (2005).

26. Okada, H., Ebhardt, H. A., Vonesch, S. C., Aebersold, R. & Hafen, E. Proteome-wide association studies identify biochemical modules associated with a wing-size phenotype in *Drosophila melanogaster*. Nat Commun 7, 12649, doi:10.1038/ncomms12649 (2016).

27. Tregenza, T. & Wedell, N. Definitive evidence for cuticular pheromones in a cricket. Anim Behav 54, 979–984 (1997).

28. Thomas, M. L. & Simmons, L. W. Sexual selection on cuticular hydrocarbons in the Australian field cricket, *Teleogryllus oceanicus*. BMC Evol Biol 9, 162, doi:10.1186/1471-2148-9-162 (2009).

29. Thomas, M. L. & Simmons, L. W. Cuticular hydrocarbons influence female attractiveness to males in the Australian field cricket, *Teleogryllus oceanicus*. J Evol Biol 23, 707–714 (2010).

30. Simmons, L. W., Thomas, M. L., Gray, B. & Zuk, M. Replicated evolutionary divergence in the cuticular hydrocarbon profile of male crickets associated with the loss of song in the Hawaiian archipelago. J Evol Biol 27, 2249–2257 (2014).

31. Gray, B., Bailey, N. W., Poon, M. & Zuk, M. Multimodal signal compensation: do field crickets shift sexual signal modality after the loss of acoustic communication? Animal behaviour 93, 243–248 (2014).

32. Dugatkin, L. A. Principles of Animal Behavior. 2nd edn, (W. W. Norton & Company, 2008).

33. Bank, C., Hietpas, R. T., Wong, A., Bolon, D. N. & Jensen, J. D. A bayesian MCMC approach to assess the complete distribution of fitness effects of new mutations: uncovering the potential for adaptive walks in challenging environments. Genetics 196, 841–852 (2014).

34. Pascoal, S. et al. Rapid evolution and gene expression: a rapidly evolving Mendelian trait that silences field crickets has widespread effects on mRNA and protein expression. J Evol Biol 29, 1234–1246 (2016).

35. Bailey, N. W., Gray, B. & Zuk, M. Acoustic experience shapes alternative mating tactics and reproductive investment in male field crickets. Curr Biol 20, 845–849 (2010).

36. Pascoal, S. et al. Increased socially mediated plasticity in gene expression accompanies rapid adaptive evolution. Ecol Lett 21, 546–556 (2018).

37. Kupper, C. et al. A supergene determines highly divergent male reproductive morphs in the ruff. Nat Genet 48, 79–83 (2016).

38. Joag, R. et al. Transcriptomics of intralocus sexual conflict: Gene expression patterns in females change in response to selection on a male secondary sexual trait in the bulb mite. Genome Biol Evol 8, 2351–2357 (2016).

39. Martin, M. Cutadapt removes adapter sequences from high-throughput sequencing reads. EMBnet. journal 17, pp. 10–12 (2011).

40. Heo, Y., Wu, X. L., Chen, D., Ma, J. & Hwu, W. M. BLESS: bloom filter-based error correction solution for high-throughput sequencing reads. Bioinformatics 30, 1354–1362 (2014).

41. Rognes, T., Flouri, T., Nichols, B., Quince, C. & Mahe, F. VSEARCH: a versatile open source tool for metagenomics. PeerJ 4, e2584, doi:10.7717/peerj.2584 (2016).

42. Kajitani, R. et al. Efficient de novo assembly of highly heterozygous genomes from whole-genome shotgun short reads. Genome Res 24, 1384–1395 (2014).

43. English, A. C. et al. Mind the gap: upgrading genomes with Pacific Biosciences RS long-read sequencing technology. PLoS One 7, e47768, doi:10.1371/journal.pone.0047768 (2012).

44. Walker, B. J. et al. Pilon: an integrated tool for comprehensive microbial variant detection and genome assembly improvement. PLoS One 9, e112963, doi:10.1371/journal.pone.0112963 (2014).

45. Simao, F. A., Waterhouse, R. M., Ioannidis, P., Kriventseva, E. V. & Zdobnov, E. M. BUSCO: assessing genome assembly and annotation completeness with single-copy orthologs. Bioinformatics 31, 3210–3212 (2015).

46. Tarailo-Graovac, M. & Chen, N. Using RepeatMasker to identify repetitive elements in genomic sequences. Curr Protoc Bioinformatics Chapter 4, Unit 4 10, doi:10.1002/0471250953.bi0410s25 (2009).

47. Price, A. L., Jones, N. C. & Pevzner, P. A. De novo identification of repeat families in large genomes. Bioinformatics 21 Suppl 1, i351–358, doi:21/suppl_1/i351 (2005).

48. Hubley, R. et al. The Dfam database of repetitive DNA families. Nucleic Acids Res 44, D81–89, doi:10.1093/nar/gkv1272(2016).

49. Bao, W., Kojima, K. K. & Kohany, O. Repbase Update, a database of repetitive elements in eukaryotic genomes. Mob DNA 6, 11, doi:10.1186/s13100-015-0041-9 (2015).

50. Boratyn, G. M. et al. Domain enhanced lookup time accelerated BLAST. Biol Direct 7, 12, doi:10.1186/1745-6150-7-12 (2012).

51. Smit, A., Hubley, R. & Green, P. RepeatMasker Open-4.0, <http://www.repeatmasker.org> (2013-2015).

52. Abrusan, G., Grundmann, N., DeMester, L. & Makalowski, W. TEclass--a tool for automated classification of unknown eukaryotic transposable elements. Bioinformatics 25, 1329–1330 (2009).

53. Benson, G. Tandem repeats finder: a program to analyze DNA sequences. Nucleic Acids Res 27, 573–580 (1999).

54. Korf, I. Gene finding in novel genomes. BMC Bioinformatics 5, 59, doi:10.1186/1471-2105-5-59 (2004).

55. Majoros, W. H., Pertea, M. & Salzberg, S. L. TigrScan and GlimmerHMM: two open source ab initio eukaryotic gene-finders. Bioinformatics 20, 2878–2879 (2004).

56. Blanco, E., Parra, G. & Guigo, R. Using geneid to identify genes. Curr Protoc Bioinformatics Chapter 4, Unit 4 3, doi:10.1002/0471250953.bi0403s18 (2007).

57. Hoff, K. J., Lange, S., Lomsadze, A., Borodovsky, M. & Stanke, M. BRAKER1: Unsupervised RNA-Seq-Based Genome Annotation with GeneMark-ET and AUGUSTUS. Bioinformatics 32, 767–769 (2016).

58. Kim, D. et al. TopHat2: accurate alignment of transcriptomes in the presence of insertions, deletions and gene fusions. Genome Biol 14, R36, doi:10.1186/gb-2013-14-4-r36 (2013).

59. Li, H. et al. The Sequence Alignment/Map format and SAMtools. Bioinformatics 25, 2078–2079 (2009).

60. Lomsadze, A., Burns, P. D. & Borodovsky, M. Integration of mapped RNA-Seq reads into automatic training of eukaryotic gene finding algorithm. Nucleic Acids Res 42, e119, doi:10.1093/nar/gku557 (2014).

61. Stanke, M., Diekhans, M., Baertsch, R. & Haussler, D. Using native and syntenically mapped cDNA alignments to improve de novo gene finding. Bioinformatics 24, 637–644 (2008).

62. Wang, X. et al. The locust genome provides insight into swarm formation and long-distance flight. Nat Commun 5, 2957,, doi:10.1038/ncomms3957 (2014).

63. The International Aphid Genomics Consortium. Genome sequence of the pea aphid *Acyrthosiphon pisum*. PLoS Biol 8, e1000313, doi:10.1371/journal.pbio.1000313 (2010).

64. Gramates, L. S. et al. FlyBase at 25: looking to the future. Nucleic Acids Res 45, D663–D671, doi:10.1093/nar/gkw1016 (2017).

65. McKenna, D. D. et al. Genome of the Asian longhorned beetle (*Anoplophora glabripennis*), a globally significant invasive species, reveals key functional and evolutionary innovations at the beetle-plant interface. Genome Biol 17, 227, doi:10.1186/s13059-016-1088-8 (2016).

66. Xue, J. et al. Genomes of the rice pest brown planthopper and its endosymbionts reveal complex complementary contributions for host adaptation. Genome Biol 15, 521, doi:10.1186/s13059-014-0521-0 (2014).

67. Benoit, J. B. et al. Unique features of a global human ectoparasite identified through sequencing of the bed bug genome. Nat Commun 7, 10165, doi:10.1038/ncomms10165 (2016).

68. Suyama, M., Torrents, D. & Bork, P. BLAST2GENE: a comprehensive conversion of BLAST output into independent genes and gene fragments. Bioinformatics 20, 1968–1970 (2004).

69. Birney, E., Clamp, M. & Durbin, R. GeneWise and Genomewise. Genome Res 14, 988–995 (2004).

70. Grabherr, M. G. et al. Full-length transcriptome assembly from RNA-Seq data without a reference genome. Nat Biotechnol 29, 644–652 (2011).

71. Xu, Y., Wang, X., Yang, J., Vaynberg, J. & Qin, J. PASA--a program for automated protein NMR backbone signal assignment by pattern-filtering approach. J Biomol NMR 34, 41–56 (2006).

72. Haas, B. J. et al. Automated eukaryotic gene structure annotation using EVidenceModeler and the Program to Assemble Spliced Alignments. Genome Biol 9, R7, doi:10.1186/gb-2008-9-1-r7 (2008).

73. Yang, Y. et al. Draft genome of the Marco Polo Sheep (*Ovis ammon polii*). GigaScience 6, 1–7, doi:10.1093/gigascience/gix106 (2017).

74. Finn, R. D. et al. InterPro in 2017-beyond protein family and domain annotations. Nucleic Acids Res 45, D190–D199, doi:10.1093/nar/gkw1107 (2017).

75. Bairoch, A. & Apweiler, R. The SWISS-PROT protein sequence database and its supplement TrEMBL in 2000. Nucleic Acids Res 28, 45–48 (2000).

76. Jones, P. et al. InterProScan 5: genome-scale protein function classification. Bioinformatics 30, 1236–1240 (2014).

77. The Gene Ontology Consortium. Expansion of the Gene Ontology knowledgebase and resources. Nucleic Acids Res 45, D331–D338, doi:10.1093/nar/gkw1108 (2017).

78. Adams, M. D. et al. The genome sequence of *Drosophila melanogaster*. Science 287, 2185–2195 (2000).

79. Gotz, S. et al. High-throughput functional annotation and data mining with the Blast2GO suite. Nucleic Acids Res 36, 3420–3435 (2008).

80. Kanehisa, M., Sato, Y. & Morishima, K. BlastKOALA and GhostKOALA: KEGG tools for functional characterization of genome and metagenome sequences. J Mol Biol 428, 726–731 (2016).

81. Tang, H. et al. ALLMAPS: robust scaffold ordering based on multiple maps. Genome Biol 16, 3, doi:10.1186/s13059-014-0573-1 (2015).

82. Kent, W. J. et al. The human genome browser at UCSC. Genome Res 12, 996–1006 (2002).

83. Pascoal, S. et al. Sexual selection and population divergence I: The influence of socially flexible cuticular hydrocarbon expression in male field crickets (*Teleogryllus oceanicus*). Evolution 70, 82–97 (2016).

84. Catchen, J., Hohenlohe, P. A., Bassham, S., Amores, A. & Cresko, W. A. Stacks: an analysis tool set for population genomics. Mol Ecol 22, 3124–3140 (2013).

85. Li, H. & Durbin, R. Fast and accurate short read alignment with Burrows-Wheeler transform. Bioinformatics 25, 1754–1760 (2009).

86. Calus, M. P. Genomic breeding value prediction: methods and procedures. Animal 4, 157–164 (2010).

87. VanRaden, P. M. Efficient methods to compute genomic predictions. J Dairy Sci 91, 4414–4423 (2008).

88. Donoughe, S. & Extavour, C. G. Embryonic development of the cricket *Gryllus bimaculatus*. Dev Biol 411, 140–156 (2016).

89. Joshi, N. A. & Fass, J. N. SICKLE: a sliding-window, adaptive, quality-based trimming tool for FastQ files [Software], <https://github.com/najoshi/sickle> (2011).

90. Pertea, M., Kim, D., Pertea, G. M., Leek, J. T. & Salzberg, S. L. Transcript-level expression analysis of RNA-seq experiments with HISAT, StringTie and Ballgown. Nat Protoc 11, 1650–1667 (2016).

91. Robinson, M. D., McCarthy, D. J. & Smyth, G. K. edgeR: a Bioconductor package for differential expression analysis of digital gene expression data. Bioinformatics 26, 139–140 (2010).

92. Kanehisa, M., Sato, Y., Kawashima, M., Furumichi, M. & Tanabe, M. KEGG as a reference resource for gene and protein annotation. Nucleic Acids Res 44, D457–462, doi:10.1093/ nar/gkv1070 (2016).

93. Schneider, W. T., Rutz, C., Hedwig, B. & Bailey, N. W. Vestigial singing behaviour persists after the evolutionary loss of song in crickets. Biol Lett 14, doi:10.1098/rsbl.2017.0654 (2018).

94. Li, S. et al. The genomic and functional landscapes of developmental plasticity in the American cockroach. Nat Commun 9, 1008, doi:10.1038/s41467-018-03281-1 (2018).

95. Rosenfeld, J. A. et al. Genome assembly and geospatial phylogenomics of the bed bug *Cimex lectularius*. Nat Commun 7, 10164, doi:10.1038/ncomms10164 (2016).

96. Terrapon, N. et al. Molecular traces of alternative social organization in a termite genome. Nat Commun 5, 3636, doi:10.1038/ncomms4636 (2014).

97. Dudchenko, O. et al. De novo assembly of the *Aedes aegypti* genome using Hi-C yields chromosome-length scaffolds. Science 356, 92–95 (2017).

98. Miller, J. R. et al. Analysis of the *Aedes albopictus* C6/36 genome provides insight into cell line utility for viral propagation. GigaScience 7, 1–13, doi:10.1093/gigascience/gix135 (2018).

99. Majlat, P., Erdos, Z. & Takacs, J. Calculation and application of retention indexes in programmed-temperature gas-chromatography. J Chromatogr 91, 89–103 (1974).

